# TSC2-extracellular matrix crosstalk controls pulmonary vascular proliferation and pulmonary hypertension

**DOI:** 10.1101/2021.08.10.455847

**Authors:** Yuanjun Shen, Dmitry A. Goncharov, Andressa Pena, Jeffery Baust, Andres Chavez Barragan, Arnab Ray, Analise Rode, Timothy N. Bachman, Baojun Chang, Mauricio Rojas, Horace DeLisser, Ana L. Mora, Tatiana V. Kudryashova, Elena. A. Goncharova

## Abstract

Increased proliferation and survival of resident cells in small pulmonary arteries (PA) are important drivers of pulmonary hypertension (PH). Tuberous sclerosis complex 2 (TSC2) is a negative regulator of mTOR complex 1 and cell growth. Here we show that TSC2 is deficient in small remodeled PA/PA vascular smooth muscle cells (PAVSMC) from human PAH and experimental PH lungs. TSC2 deficiency was reproduced *in vitro* by maintaining PAVSMC on pathologically stiff substrates and was required for stiffness-induced proliferation, accumulation of transcriptional co-activators YAP/TAZ and up-regulation of mTOR. Depletion of TSC2 reproduced PH features *in vitro* in human PAVSMC and *in vivo* in SM22-Tsc2+/− mice. TSC2 loss in PAVSMC was supported by YAP and led to the up-regulation of YAP/TAZ and mTOR via modulating the extracellular matrix (ECM) composition. ECM, produced by TSC2-deficient PAVSMC, promoted growth of non-diseased PA adventitial fibroblasts and PAVSMC, which, in turn, was prevented by α5β1 integrin receptor antagonist ATN161. *In vitro*, molecular and pharmacological (SRT2104) restoration of TSC2 down-regulated YAP/TAZ, mTOR, and ECM production, inhibited proliferation and induced apoptosis in human PAH PAVSMC. *In vivo*, orally administrated SRT2104 restored TSC2, resolved pulmonary vascular remodeling, PH, and improved right heart in two rodent models of PH. Thus, PAVSMC TSC2 is a critical integrator of ECM composition and stiffness with pro-proliferative signaling and PH, and the restoration of functional TSC2 could be an attractive therapeutic option to treat PH.

**One Sentence Summary:** TSC2 acts as mechanosensor and mechanotransducer, integrating ECM composition and stiffness with pro-proliferative signaling in pulmonary vasculature; its deficiency in PA vascular smooth muscle cells results in ECM remodeling, hyper-proliferation and pulmonary hypertension, which could be reversed by pharmacological restoration of functional TSC2.

## Introduction

Pulmonary arterial hypertension (PAH), a progressive and rapidly fatal disease with a high mortality rate and no curative options (1), represents a serious public health problem with continuously increasing death and hospitalization rates (2–4). PAH manifests by vasoconstriction and remodeling of small pulmonary arteries (PA) (5–10) leading to chronically increased PA pressure and right ventricular afterload and resulting in right heart failure and death (4). Available therapies fail to reverse established pulmonary vascular remodeling or prevent disease progression (5, 9), and development of novel anti-proliferative/anti-remodeling therapies is an area of unmet important need.

Increased proliferation and survival of resident pulmonary vascular cells, a key components of PA remodeling (8, 11, 12), are induced by the combination of soluble and insoluble stimuli, such as excessive production of growth factors and pro-inflammatory mediators and stiffening and remodeling of the extracellular matrix (ECM). As PAH progresses, PA vascular smooth muscle cells (PAVSMC) undergo a switch to the secretory, proliferative, apoptosis-resistant phenotype, which is self-supported by constitutive inhibition of growth suppressor cassette HIPPO-LATS1 and consequent up-regulation of its pro-proliferative downstream effectors transcriptional co-activators Yes-associated protein (YAP)/TAZ and the mechanistic target of rapamycin (mTOR) complex 1 (mTORC1) and mTORC2 (7–9, 12–15). The mechanisms coordinating growth-promoting signals with proliferative responses of resident PA cells in PAH are not completely understood, and available anti-proliferative therapeutic strategies are challenging due to their side effects.

Growth suppressor tuberous sclerosis complex 2 (TSC2) (tuberin) is a negative regulator of mTORC1, cell growth and proliferation (16). Deficiency or mutational inactivation of TSC2 is linked to proliferative diseases such as cancer, tuberous sclerosis and pulmonary lymphangioleiomyomatosis (LAM) (16–18). Several lines of evidence suggest that TSC2 may act as a coordinator of mTORC1 and HIPPO-YAP/TAZ pathways in PAH pulmonary vasculature. First, mTORC1 is activated in small remodeled PAs and contributes to human PAH PAVSMC proliferation, VSM remodeling and experimental PH in mice (7, 19, 20). Second, mTOR-induced accumulation of YAP has been reported in Tsc2-null cells (21). Next, mice with VSM-specific knock-down of Tsc1 (a binding partner of TSC2 protecting it from the degradation) develop PH (22). Lastly, a subset of patients with pulmonary LAM (caused by TSC somatic mutations or loss of heterozygosity) develop “out-of-proportion” pre-capillary PH, which can’t be fully explained by hypoxemia and is likely attributed to the loss of pulmonary vascular TSC function (23).

In the present study, we aimed to determine the role of TSC2 in pulmonary vascular hyper-proliferation in PAH. We found that TSC2 acts as a mechanosensor and mechanotransducer, and its deficiency in smooth muscle cells from small PAs permits up-regulation of YAP-TAZ and mTOR, increases PAVSMC proliferation, survival, and remodeling, and promotes PH. We also provide a novel mechanistic link from PAVSMC-specific TSC2 deficiency to excessive ECM production, ECM-dependent activation of YAP/TAZ and mTOR, and increased growth of PAVSMC and PA adventitial fibroblasts (PAAF). Lastly, we demonstrate benefits of TSC2 restoration by SRT2104 to inhibit YAP/TAZ and mTOR axes, reverse pulmonary vascular remodeling, and reduce PH.

## Results

### TCS2 is deficient in PAVSMC from human PAH lungs

To determine the status of TSC2 in human PAH lungs, we first performed immunohistochemical and immunoblot analysis of lung tissue specimens from subjects with PAH and non-diseased (control) donor lungs (see **Table S1** for human subjects’ characteristics). We found that TSC2 protein levels are markedly lower in smooth muscle α-actin (SMA)-positive areas and whole tissue lysates of small PAs from PAH lungs compared to controls (**Fig. 1a-d**). Supporting our observations, PAVSMC from small (<1.5 mm outer diameter) human PAH PAs had significantly reduced TSC2 protein content (**Fig. 1e, f**) and significantly higher non-stimulated growth and proliferation than cells from non-diseased (control) subjects (**Fig. 1g, h**). We did not observe TSC2 deficiency in human PAH PA endothelial cells (PAEC) and PAAF (**Fig. S1**). Together, these data demonstrate that TSC2 is deficient in PAVSMC in small remodeled PAs in human PAH lungs.

**Figure 1.**
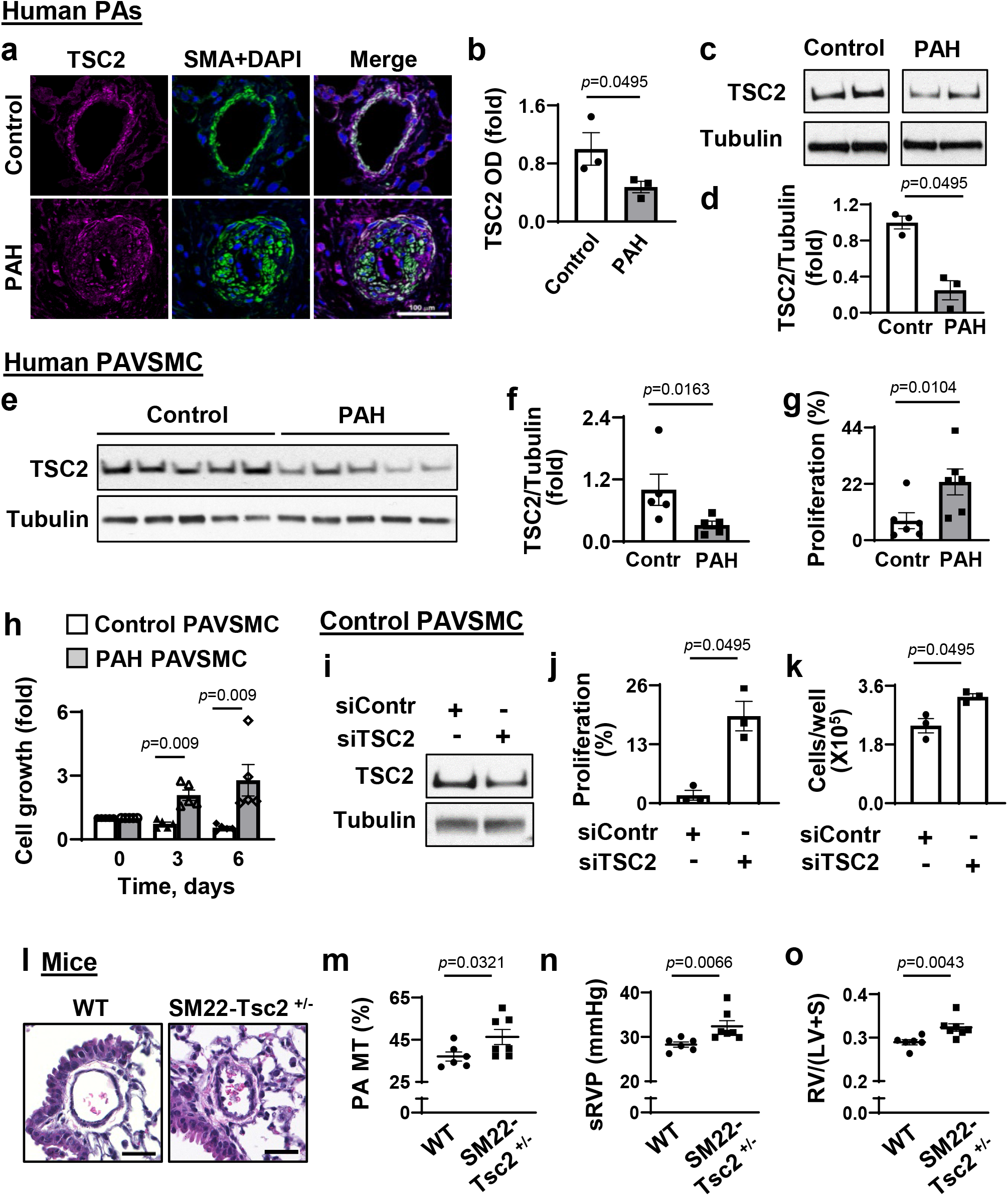
TSC2 deficiency results in the increased PAVSMC proliferation, vascular remodeling and PH. (see also Supplemental Figs. S1 and S2 for additional data) **a, b.** (**a**) Immunohistochemical analysis of human lung tissue sections to detect TSC2 (magenta), SMA (green) and DAPI (blue). Bar equals 100 μm. Images are representative from 3 subjects/group, 6 PA/subject. (**b**) Optical density (OD) measurement of TSC2 fluorescent signal in SMA-positive regions of human small PAs. Data are means±SE from 3 subjects/group, 6 PA/subject, 12 areas/PA, fold change to control. Statistical analysis by Mann Whitney U test (significance *p*<0.05), PAH vs. control. **c,d.** Immunoblot analysis of small PAs. Data are means±SE from n=3 subjects/group. Statistical analysis by Mann Whitney U test (significance *p*<0.05), PAH vs. control. **e,f.** Immunoblot analysis of human non-diseased (control) (white bars) and PAH (grey bars) PAVSMC. Data are means±SE, fold change to control; n=5 subjects/group. Statistical analysis by Mann Whitney U test (significance *p*<0.05), PAH vs. control. **g.** Proliferation (Ki67) of human control and PAH PAVSMC. Data represent % of Ki67-positive cells per total number of cells (detected by DAPI); data are means±SE from n=6 subjects/group; statistical analysis by Mann-Whitney U test (significance p<0.05), PAH vs. control. **h.** Equal quantity of cells was plated on each well of 6-well plate (day 0), cell counts were performed at days 3 and 6. Data are fold change to day 0. Data are means±SE from n=5 subjects/group; statistical analysis by Mann-Whitney U test (significance *p*<0.05), PAH vs. control. **i-k.** Immunoblot (**i**), proliferation ( DNA synthesis by BrdU incorporation) (**j**) and cell growth (cell counts) analyses (**k**) performed on the control human PAVSMC transfected with siRNA TSC2 (siTSC2) or control siRNA GLO (siContr) for 48 hours. Data are means±SE, from n=3 subjects/group; statistical analysis by Mann-Whitney U test (significance *p*<0.05), siTSC2 vs. siContr. **l-o.** Morphological and hemodynamic analysis of 9 weeks-old male SM22-Tsc2^+/−^ and wild type (WT) mice. **l:** Images are representative from 6 mice/group, 12 PA/mouse. Bar equals 30 μm. **m-o**: PA medial thickness (PA MT) (**m**), systolic RV pressure (sRVP) (**n**), and Fulton index (right ventricle (RV)/(left ventricle (LV) + septum) (**o**). Data are means±SE; n=6 mice/WT; n=7 mice/SM22-Tsc2^+/−^ group; statistical analysis by Mann-Whitney U test (significance *p*<0.05), SM22-Tsc2^+/−^ vs. WT.

### TSC2 deficiency enables PAVSMC hyper-proliferation, remodeling, and PH

To determine the functional significance of TSC2 deficiency, we first performed siRNA-induced TSC2 depletion in human control PAVSMC. Transfection with siTSC2 significantly increased DNA synthesis (BrdU incorporation) and cell growth compared to the cells transfected with control siRNA (**Fig. 1i-k**). To test *in vivo* consequences of TSC2 deficiency, we next developed mice with SMC-specific Tsc2 knockout by crossing Tsc2^flox/flox^ mice and Tagln (Sm22)-cre mice (13). Homozygous knockout of Tsc2 was embryonically lethal. Mice with heterozygous deletion of Tsc2 (Tsc2^+/−^) (**Fig. S2**) developed spontaneous mild PH as early as at nine weeks of age. Compared to the same-age wild type controls (WT), male SM22-Tsc2^+/−^ mice had significantly higher medial thickness of small (<150 μm) PAs (PA MT) (**Fig. 1l, m**), elevated systolic right ventricular (RV) pressure (sRVP) (**Fig. 1n**), and RV hypertrophy (Fulton index) (**Fig. 1o**). Four out of six female mice also developed mild PH (sRVP≥30 mmHg). Together, these data demontrate that TSC2 deficiency results in increased PAVSMC proliferation, VSM remodeling, and spontaneous PH.

### TSC2 loss is induced by substrate stiffening, enables YAP/TAZ accumulation and PAVSMC growth

Hyper-proliferation of PAVSMC in PAH is induced by multiple soluble (growth factors, inflammatory mediators) and insoluble factors (ECM composition and stiffness) (9, 11). To identify pro-PAH stimuli that induce TSC2 deficiency in PAVSMC, we exposed control human PAVSMC to the soluble growth factors and pro-inflammatory mediators (platelet-derived growth factor-BB (PDGF-BB), insulin growth factor 1 (IGF-1), interleukin-6 (IL-6), tumor necrosis factor-α (TNF-α) and endotelin-1 (ET-1), or maintained the cells on the matrices of a different composition (collagen 1, collagen IV, fibronectin, laminin) or stiffness (physiological 0.2 kPa and pathological 25 kPa) (8, 11, 13). We found that TSC2 protein levels were moderately decreased in PAVSMC stimulated by PDGF-BB or maintained on the plates covered with collagen I (**Fig. 2a-d**). Importantly, maintenance of PAVSMC on the pathologically stiff (25 kPa) (8, 12, 14, 15) matrices dramatically reduced TSC2 protein content and significantly increased cell growth compared to the cells seated on physiologically soft (0.2 kPa) substrates (**Fig. 2e-g**). To determine whether TSC2 modulates stiffness-induced PAVSMC growth, we transfected control human PAVSMC seeded on the stiff (25 kPa) matrices with mammalian vectors expressing GFP-TSC2 or control GFP. GFP-TSC2-transfected PAVSMC had ~2 times lower growth on stiff matrices than GFP-transfected cells (**Fig. 2h**), suggesting that TSC2 acts as a mechanosensor and mechanotransducer, and TSC2 deficiency in PAVSMC is induced by increased matrix stiffness and is required for stiffness-induced cell growth.

**Figure 2.**
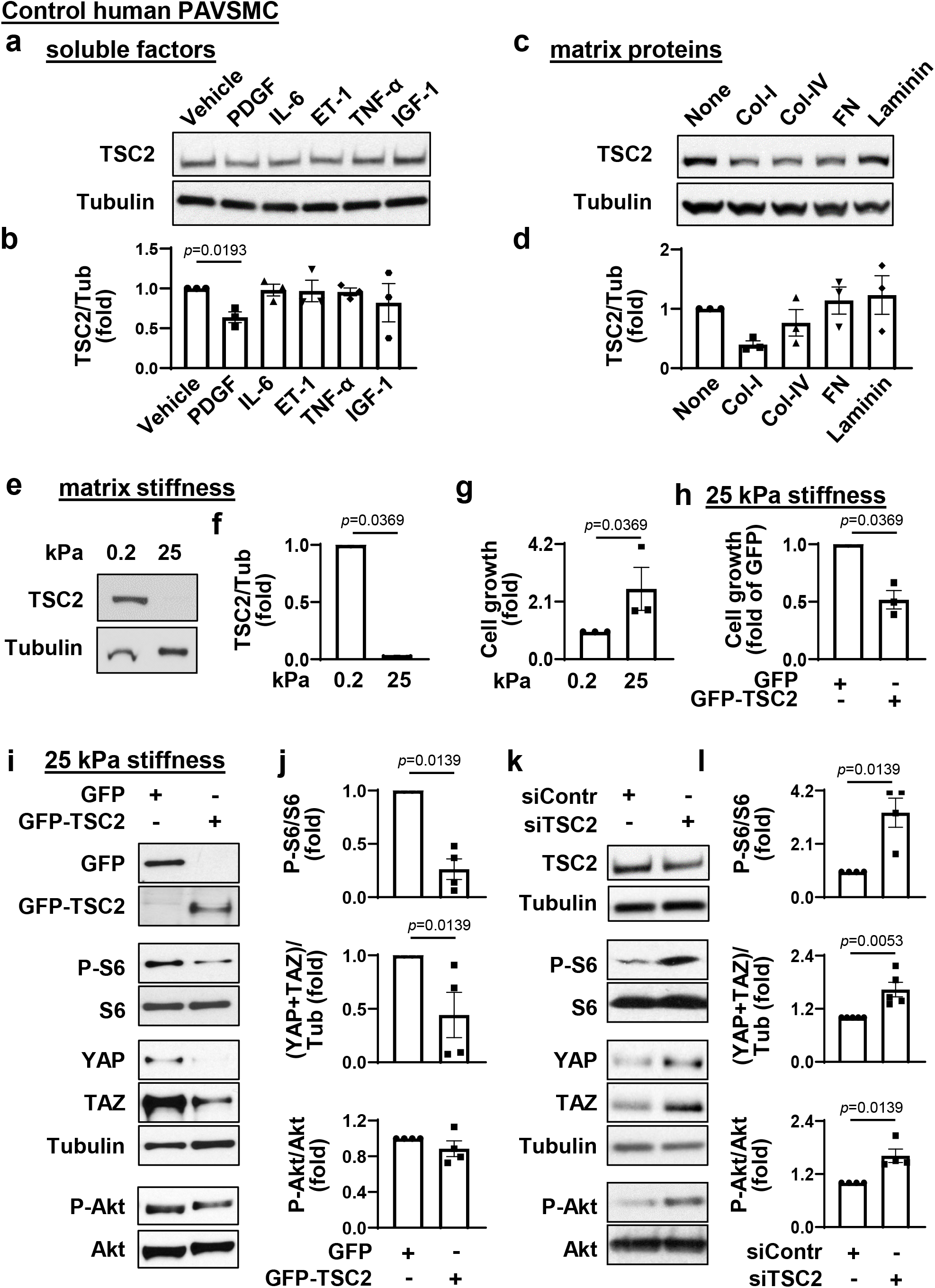
TSC2 deficiency is induced by increased substrate stiffness and is required for YAP/TAZ accumulation, activation of Akt and mTOR, and increased growth of PAVSMC. **a-d.** Immunoblot analysis of control human PAVSMC treated with indicated soluble factors (**a,b**) or maintained on the indicated matrices (**c,d**) for 48 hours. Data are means±SE; fold change to vehicle treatment or none-coated matrix from n=3 subjects/group; statistical analysis by Kruskal-Wallis test with Dunn’s Pairwise Comparison (significance *p*<0.025). **e, f.** Immunoblot analysis of control human PAVSMC maintained on the softwell hydrogels with normal (0.2 kPa) or pathological (25 kPa) stiffness for 48 hours. Data are means±SE, fold change to 0.2 kPa from n=3 subjects/group; statistical analysis by Mann Whitney U test (significance *p*<0.05), 25 kPa vs. 0.2 kPa. **g.** Equal amounts of control human PAVSMC were plated at the hydrogels with 0.2 kPa or 25 kPa stiffness, and cell growth assay (cell counts) was performed at day 4. Data are means±SE, fold change to 0.2 kPa from n=3 subjects/group; statistical analysis by Mann Whitney U test (significance *p*<0.05), 25 kPa vs. 0.2 kPa. **h, i, j.** Human control PAVSMC, plated on the softwell hydrogels with 25 kPa stiffness were transfected with mammalian vectors expressing GFP or GFP-tagged human TSC2 for 36 hours. Cell growth assay (cell counts) and immunoblot analysis were performed. Data are means±SE, fold change to GFP transfection from n=3 (cell counts) and 4 (immunoblots) subjects/group; statistical analysis by Mann Whitney U test (significance *p*<0.05), GFP-TSC2 vs. GFP. **k, l.** Human control PAVSMC were transfected with siRNA TSC2 (siTSC2) or control siRNA GLO (siContr); 48 hours later, immunoblot analysis was performed. Data are means±SE, fold change to siContr from n=4 or 5 subjects/group; statistical analysis by Mann Whitney U test (significance *p*<0.05), siTSC2 vs. siContr.

Stiffening of small PAs drives pulmonary vascular cell proliferation and remodeling in PAH via transcriptional co-activators YAP/TAZ, which act, at least in part, via up-regulating proproliferative/pro-survival Akt-mTOR signaling (8, 12, 14). Interestingly, re-expression of TSC2 in control PAVSMC maintained on the stiff matrices not only significantly decreased phosphorylation of ribosomal protein S6, a molecular signature of mTORC1 activation (**Fig. 2i, j**), but also significantly reduced YAP/TAZ protein levels compared to the GFP-expressing cells (**Fig. 2i, j**). Supporting our findings, siRNA-dependent depletion of TSC2 in control human PAVSMC resulted in a significant increase of mTORC1-specific S6 phosphorylation, accumulation of YAP/TAZ and elevated phospho-S473-Akt compared to cells transfected with control siRNA (**Fig. 2k, l**). These data show that TSC2 deficiency in PAVSMC is required for stiffness-induced PAVSMC growth and up-regulation of YAP/TAZ and mTOR.

### TSC2 loss up-regulates YAP/TAZ, mTOR and PAVSMC proliferation via ECM remodeling

The ECM remodeling is an important factor that facilitates hyper-proliferation of resident PA cells in PAH. Because YAP and TAZ facilitate resident pulmonary vascular cell proliferation in PAH via ECM remodeling (8, 11), we hypothesized that, in addition to canonical Rheb-dependent activation of mTORC1, TSC2 deficiency in PAH PAVSMC may facilitate pro-proliferative signaling via modulating ECM production. In agreement with published studies (8, 11), PAVSMC from human PAH lungs had higher fibronectin and collagen 1A protein content compared to controls (**Fig. 3a, b**). Importantly, we found that transfection with siRNA TSC2 significantly increased fibronectin and collagen 1A protein levels in control human PAVSMC (**Fig. 3c, d**), suggesting that TSC2 loss facilitates production of the two key ECM proteins involved in PAH.

**Figure 3.**
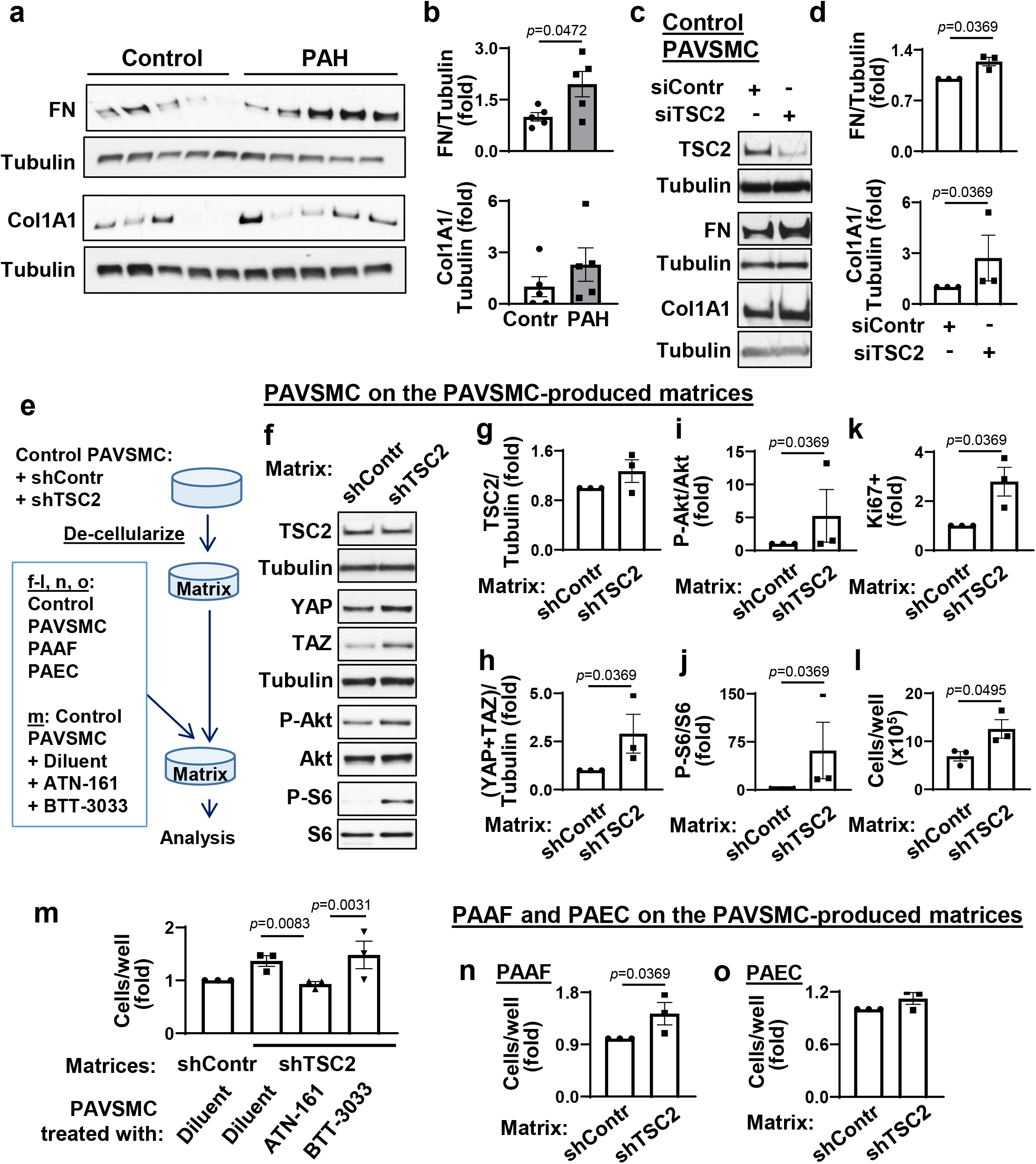
TSC2 loss in PAVSMC modulates YAP/TAZ and mTOR, promotes growth of non-diseased pulmonary vascular cells via extracellular matrix remodeling. (see also Supplemental Fig.S3, S4, and S5 for additional data) **a, b.** Immunoblot analysis of control and PAH PAVSMC (5 subjects/group) to detect fibronectin (FN) and collagen 1A1 (Col1A1). Data are means±SE, fold change to control. p<0.05 by Mann-Whitney U test. **c, d.** Immunoblot analysis of human control PAVSMC transfected with siRNA TSC2 (siTSC2) or control siRNA GLO (siContr) to detect FN and Col1A1. Data are means±SE from n=3 subjects/group, fold change to siContr; statistical analysis by Mann Whitney U test (significance *p*<0.05). **e-o**. Pre-confluent control PAVSMC were infected with shRNA control (shContr) or shRNA TSC2 (shTSC2) adenovirus and maintained for 6 days. Then cells were removed and equal amount of non-diseased (control) PAVSMC, untreated (**f-l**) or PAVSMC treated with diluent, 10μM ATN161 (integrin α5β1 inhibitor), or 10μM BTT3033 (integrin α2β1 inhibitor) (**m**), control PAAF (**n**), or control PAEC (**o**) were plated on the remaining matrices. Four days post-plating, immunoblot analysis (**f-j**), proliferation (Ki67) (**k**), and cell growth assays (cell counts) (**l-q**) were performed. Data are means±SE from n=3 subjects/group, statistical analysis by Mann Whitney U test (significance *p*<0.05) (**g-l, n, o**) and by Kruskal-Wallis test with Dunn’s Pairwise Comparison (significance *p*<0.025). (**m**).

Next, we evaluated whether TSC2 loss modulates pro-proliferative signaling of neighboring cells via ECM. We prepared de-cellularized matrices from control human PAVSMC infected with lentiviruses producing shTSC2 or control shRNA and used these cell-free matrices as substrates for plating of control human PAVSMC (**Fig. 3e**). We found that control PAVSMC, seeded on the matrices produced by shTSC2-infected control PAVSMC, had significantly higher YAP/TAZ protein levels, increased S473-Akt and ribosomal protein S6 phosphorylation rates, and elevated growth and proliferation compared to the cells seeded on the matrices produced by shContr-infected PAVSMC (**Fig. 3f-l**), demonstrating that TSC2-deficient PAVSMC up-regulate YAP/TAZ, Akt, mTOR, and cell proliferation via ECM. To further confirm our observations, we maintained control PAVSMC on the cell-free matrices, produced by shTSC2-infected PAVSMC, in the presence of ATN-161, a small peptide antagonist of integrin α5β1, and BTT-3033, a selective inhibitor of integrin α_2_β_1_, to block cell binding with fibronectin and collagen respectively (**Fig. 3e**) and performed a cell growth assay. Interestingly, ATN-161, but not BTT3033, prevented matrix-induced PAVSMC growth (**Fig. 3m**), suggesting that TSC2 loss promotes growth of PAVSMC via regulating extracellular fibronectin content. Supporting the relevance of these findings to human PAH, control PAVSMC seeded at the de-cellularized ECM, produced by human PAH PAVSMC, had higher S-473Akt and mTORC1-specific S6 phosphorylation rates and YAP/TAZ protein content than cells maintained on the matrices produced by non-diseased (control) PAVSMC (**Fig. S3**). Collectively, these data demonstrate that TSC2 deficiency promotes YAP/TAZ and mTOR up-regulation and PAVSMC proliferation via ECM remodeling.

Interestingly, in contrast to stiffness-induced TSC2 deficiency (**Fig. 2e, f**), control PAVSMC seeded on the “diseased” matrices produced by TSC2-deficient control or PAH PAVSMC had up-regulated YAP/TAZ and mTOR without reduction of TSC2 protein levels (**Fig. 3f-i, S3**). These data support our observations that TSC2 acts upstream of YAP/TAZ and mTOR via regulating ECM and suggest an existence of ECM composition-independent mechanism(s) maintaining self-sustaining TSC2 deficiency in PAH PAVSMC. Because mTOR acts both upstream (mTORC2) and downstream (mTORC1) of TSC2 (17), and both mTORC2 and mTORC1 are up-regulated and act downstream of YAP/TAZ in human PAH PAVSMC (7, 8), we hypothesized that stiffness-independent TSC2 deficiency could be self-supported via a YAP/mTORC2 feed-forward loop. Supporting our hypothesis, depletion of YAP with specific siRNA led to TSC2 accumulation in human PAH PAVSMC compared to cells transfected with control siRNA (**Fig. S4**). However, treatment of human PAH PAVSMC with either mTOR kinase inhibitor PP242 (which inhibits mTOR in both mTORC1 and mTORC2) or allosteric mTORC1 inhibitor rapamycin did not increase TSC2 protein levels or reduce YAP/TAZ accumulation (**Fig. S5**), suggesting that TSC2 deficiency in PAH PAVSMC is supported by YAP in the mTOR-independent manner.

### ECM produced by TSC2-deficient PAVSMC induces growth of PAAF

Because changes in ECM composition affect the proliferative response of neighboring pulmonary vascular cells, we next tested whether the matrix, produced by TSC2-deficient PAVSMC, modulates growth of other resident pulmonary vascular cells. We seeded control human PAEC and PAAF on the matrices produced by control human PAVSMC infected with adenovirus producing shTSC2 or control shRNA (shContr) (**Fig. 3e**). We found that control PAAF, but not PAEC, maintained on the matrices from shTSC2-infected control PAVSMC, had significantly higher cell growth compared to the cells seeded on the matrices produced by shContr-infected PAVSMC (**Fig. 3n, o).** These data show that TSC2 deficiency in PAVSMC results in dysregulated ECM production, which promotes growth of PAAF.

### Restoration of TSC2 suppresses proliferation, induces apoptosis in PAH PAVSMC

To evaluate potential benefits of restoring functional TSC2 to target increased ECM proteins’ production, proliferation and survival of PAVSMC from human PAH lungs, we transfected human PAH PAVSMC with mammalian vectors expressing GFP-TSC2 or control GFP. We found that TSC2 suppressed mTORC1-dependent S6 phosphorylation, and significantly reduced YAP/TAZ protein levels, Akt phosphorylation rates, and fibronectin and collagen 1A production compared to GFP-expressing cells (**Fig. 4a,b, S6a**). Importantly, expression of TSC2 suppressed PAH PAVSMC proliferation (**Fig. 4c, Fig. S6b**) and induced accumulation of Cleaved Caspase 3 and significant apoptosis (**Fig. 4d, e**), suggesting that restoration of TSC2 may be considered as an attractive therapeutic strategy to suppress both YAP/TAZ and mTOR pathways, halt excessive ECM production, inhibit proliferation, and induce apoptosis in PAH PAVSMC.

**Figure 4.**
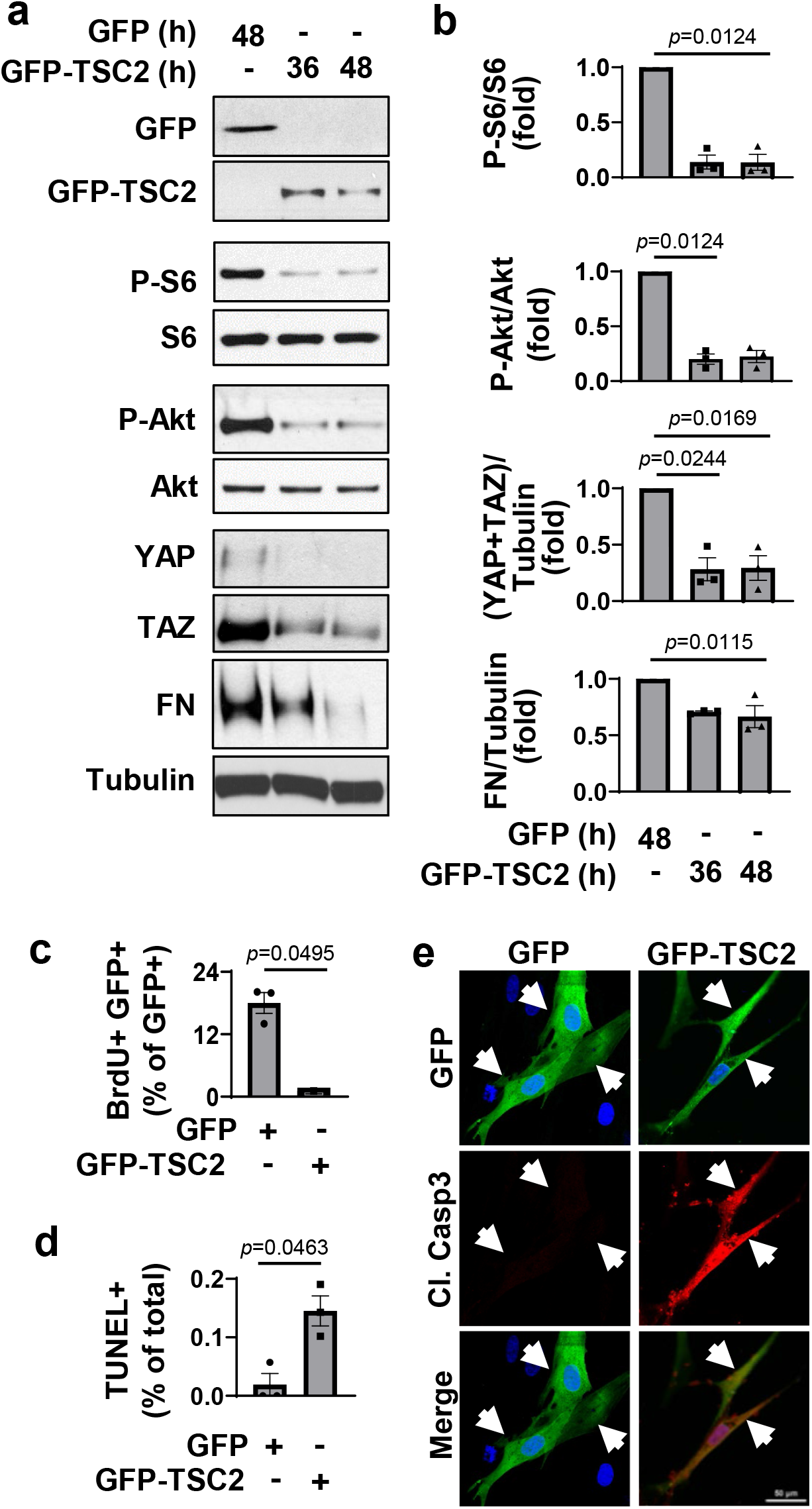
TSC2 expression reduces YAP/TAZ accumulation and fibronectin production, suppresses proliferation and induces apoptosis in human PAH PAVSMC. (see also Supplemental Fig. S6 for additional data) **a, b.** Human PAH PAVSMC were transfected with mammalian vectors expressing GFP or GFP-TSC2 for indicated times followed by immunoblot analysis to detect indicated proteins. Data are means±SE from 3 subjects/group; statistical analysis by Kruskal-Wallis rank test with Dunn’s Pairwise Comparison (significance *p*<0.025). **c-e.** DNA synthesis (BrdU) (**c**), apoptosis (TUNEL) (**d**), and immunocytochemical analysis to detect cleaved caspase 3 (Cl. Casp3, red), GFP (green), and DAPI (blue). Data are means±SE from 3 subjects/group; statistical analysis by Mann-Whitney U test (significance *p*<0.05). Images are representative from cells from 3 different subjects; a minimum of 12 transfected cells/subject/condition. Bar equals 50 μm. White arrows indicate transfected cells.

### SRT2104 reverses PAH PAVSMC phenotype via restoration of TSC2

Next, we evaluated potential pharmacological strategies to restore functional TSC2 in human PAH PAVSMC. TSC2 is regulated by direct phosphorylation, which either prevents of promotes its degradation (17), and epigenetic modifications, such as deacetylation by SIRT1 (24). Known positive and negative regulators of TSC2 involved in PAH are Akt and AMPK. Akt is activated, while AMPK is inhibited in human PAH PAVSMC, and either inhibition of Akt or activation of AMPK suppresses mTORC1 and PAH PAVSMC proliferation (8, 19, 25–28). Intriguingly, we found that the inhibition of Akt or activation of AMPK, while successfully reducing mTORC1-dependent S6 phosphorylation, did not increase TSC2 protein levels in human PAH PAVSMC (**Fig. S7**). Together with an inability of mTORC1/2 inhibitor PP242 to restore TSC2 in PAH PAVSMC (**Fig. S5**), our data suggest that mTORC2, Akt and AMPK act downstream of or in parallel to TSC2, and its targeting couldn’t restore PAH-specific TSC2 deficiency.

Next, we tested the effect of SIRT1 activator SRT2104. We found that treatment of human PAH PAVSMC with SRT2104 significantly increased TSC2 protein levels and suppressed mTORC1-dependent S6 phosphorylation (**Fig. 5a, b, e**), showing that SRT2104 restores functional TSC2. Importantly, SRT2104 reduced YAP/TAZ accumulation, decreased S473Akt phosphorylation rates, collagen 1A and fibronectin protein levels (**Fig. 5a, c, d, f, g, h**), inhibited proliferation, and promoted apoptosis in PAH PAVSMC (**Fig. 5i-k**). Importantly, SRT2104 also slowed down growth of control PAVSMC on the pathologically stiff matrices (**Fig. 5l**), suggesting potential benefits of SRT2104 in targeting both, self-sustaining and stiffness-induced PAVSMC proliferation in PAH.

**Figure 5.**
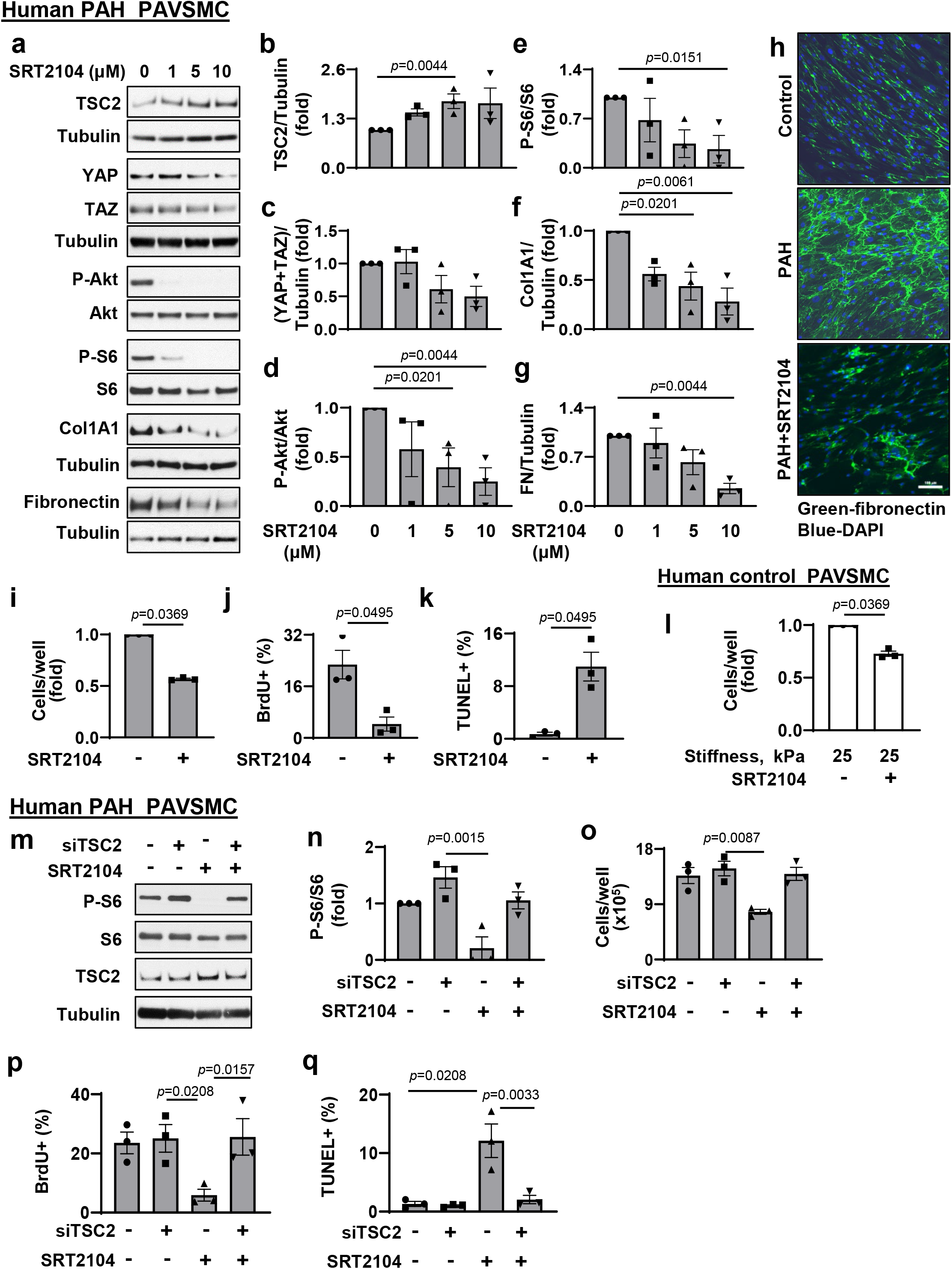
SRT2104 restores functional TSC2, inhibits proliferation and induces apoptosis in human PAH PAVSMC. (see also Supplemental Fig. S7 for additional data) **a-g.** Immunoblot analysis of human PAH PAVSMC treated with diluent (0) or indicated concentrations of SRT2104 to detect indicated proteins. Data are means±SE from 3 experiments, each performed on the cells from different subject; statistical analysis by Kruskal-Wallis rank test with Dunn’s Pairwise Comparison (significance *p*<0.025), vs. diluent (0). **h.** Immunocytochemical analysis to detect fibronectin of human non-diseased (control) and PAH PAVSMC treated with diluent or 10 μM SRT2104 for 48 hr. Bar equals 100 μm. **i-k.** Human PAH PAVSMC were treated with diluent (−) μM SRT2104 (+) for 48 hr; cell counts (**i**), proliferation (BrdU incorporation) (**j**) and apoptosis (TUNEL) (**k**) analyses were performed. Data are means±SE from 3 experiments, each performed on the cells from different subject; statistical analysis by Mann-Whitney U test (significance *p*<0.05) vs. diluent (0). **l**. Equal amount of human control PAVSMC were seeded on hydrogels with 25 kPa stiffness and treated with diluent (−) or 10 μM SRT2104 (+) for 48hr; then cell growth analysis (cell counts) was performed. Data are means±SE from 3 experiments, each performed on the cells from different subject; statistical analysis by Mann Whitney U test (significance *p*<0.05) vs. diluent (0). **m-q.** Immunoblot, cell growth, proliferation (BrdU incorporation) and apoptosis (TUNEL) analyses of PAH PAVSMC transfected with siContr or siTSC2 and treated with 10μM SRT2104 or diluent for 48hr. Data are means±SE from 3 experiments, each performed on the cells from different subject; statistical analysis by Kruskal-Wallis rank test with Dunn’s Pairwise Comparison (significance *p*<0.025) vs. siContr (Scr)+diluent (−/−).

To test whether SRT2104 acts via TSC2, we performed a “rescue” experiment by transfecting human PAH PAVSMC with siRNA TSC2 or control siRNA with and without SRT2104 treatment. We found that siRNA TSC2 prevented SRT2104-induced inhibition of the TSC2 down-stream effector P-S6, reversed SRT2104-induced inhibition of cell growth and proliferation, and protected cells from SRT2104-induced apoptosis (**Fig.5 m-q**). In aggregate, the data show that SRT2104 acts via targeting TSC2 and that restoration of TSC2 by SRT2104 inhibits proliferation and induces apoptosis in human PAH PAVSMC.

### SRT2104 restores Tsc2 in small PAs, reduces established PH in mice and rats

To evaluate the potential benefits of pharmacological restoration of TSC2 *in vivo*, we first examined whether short-term treatment of mice with SU5416/hypoxia (SuHx)-induced PH leads to Tsc2 accumulation in small PAs. SRT2104 and vehicle were administrated via gavage at days 15-21 after PH induction; negative controls were same-age same-gender mice maintained under normoxia (**Fig. 6a**). Like in human PAH, the vehicle-treated mice from SuHx group (SuHx+Vehicle) had reduced Tsc2 protein levels in SMA-positive areas of small PAs (**Fig. 6b**), suggesting that similar mechanisms are shared. As expected, mice in the SuHx+Vehicle group developed pulmonary vascular remodeling and PH as evidenced by significantly higher PA medial thickness (PA MT), systolic right ventricular pressure (sRVP), PA pressure (PAP), and RV hypertrophy (**Fig. 6b-f**). The SRT2104-treated mice (SuHx-SRT2104 group) had higher Tsc2 protein content in small PAs and significantly lower PA MT, sRVP, PAP, and RV hypertrophy compared to vehicle-treated SuHx mice (**Fig. 6b-f**). We observed no significant differences between male and female mice. Systolic left ventricular pressure (sLVP), mean arterial pressure (MAP), and heart rate were not significantly different between SuHx+Vehicle and SuHx+SRT2014 groups (**Fig. S8a-c**). These data show that short-term SRT2104 treatment elevates Tsc2, attenuates VSM remodeling, PH, and improves RV hypertrophy in mice with SuHx-induced PH.

**Figure 6.**
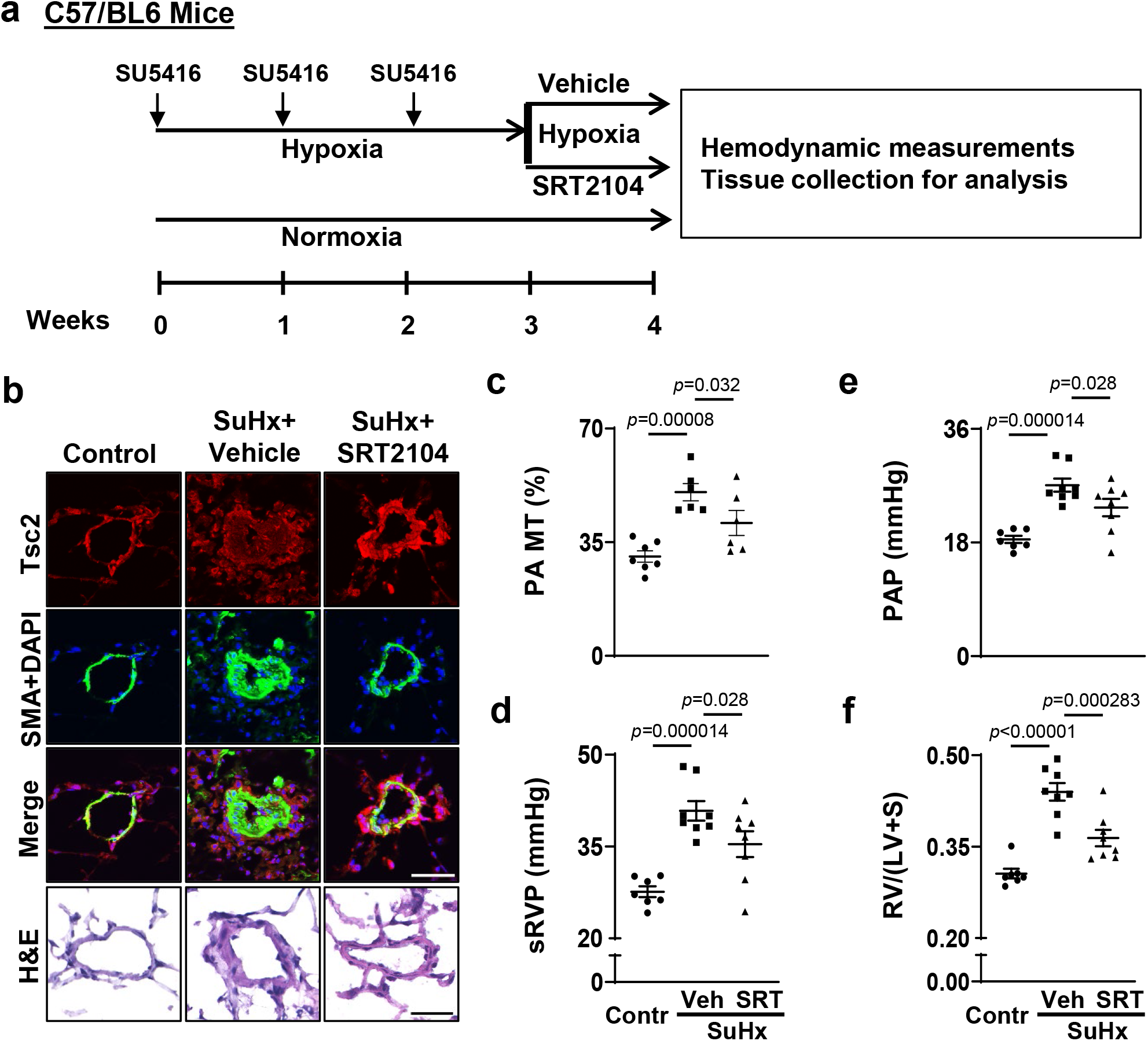
SRT2104 restores Tsc2 in small PAs, attenuates PH and reduces RV hypertrophy in mice. (see also Supplemental Fig. S8 for additional data) **a.** Schematic representation of the experiment. 6-8 weeks old male and female mice maintained under hypoxia for three weeks and received SU5416 injection at the beginning of every week. Starting week 4, mice, still kept under hypoxia, were randomly assigned to two groups and then treated with SRT2104 (SRT) or vehicle (Veh) for 1 week, 5 days/week. Controls were same-age and -sex mice kept under normoxia. Upon experiment termination, hemodynamic measurements were performed, lung and heart tissues were collected for analysis. Control mice were same-age and -sex mice maintained under normoxia. **b.** immunohistochemical (IHC) analysis to detect Tsc2 (red), SMA (green), and DAPI (blue), bar equals 80 μm; images are representative from 3 mice/group, 12 PAs/mice. H&E staining bar equals 30μm, images are representative from 7 mice/control, 6 mice/SuHx+Vehicle and SuHx+SRT2104 groups, 12 PAs/mice. **c-f. (c)** PA medial thickness (PA MT) is calculated by 12-24 PAs/mouse; data are means±SE from n=7 mice for control (3♂, 4♀), n=6 mice for PH (3♂, 3♀), n=6 mice for PH+SRT2104 (3♂, 3♀) groups; statistical analysis by one-way ANOVA (significance *p*<0.05). Systolic right ventricular pressure (sRVP) (**c**), pulmonary arterial pressure (PAP) (**d**), and Fulton index (RV/(LV + septum) weight ratio) (**e**); data are means±SE from n=7 mice for control (3♂, 4♀), n=8 mice for PH (4♂, 4♀), n=8 mice for PH+SRT2104 (4♂, 4♀) groups; statistical analysis by one-way ANOVA (significance *p*<0.05).

To evaluate potential benefits of SRT2104 in severe experimental PH, we next tested effects of long-term SRT2104 treatment using a rat SuHx model of PH. SRT5416 or vehicle were administrated to male rats for five weeks starting at day 22 of SU5416 injection (gavage, five days/week) (**Fig. 7a**). Similar to human PAH and mouse PH (**Figs. 1a, 6b**), vehicle-treated rats with SuHx-induced PH developed VSM-specific Tsc2 deficiency in small remodeled PAs (**Fig. 7b**) that was associated with robust pulmonary vascular remodeling, PH, and RV hypertrophy (**Fig. 7c-i**). Importantly, treatment with SRT2104 restored Tsc2 in small PAs, reversed pulmonary vascular remodeling, normalized sRVP and PAP, and improved RV morphology and function (reversed RV hypertrophy and normalized max dP/dT and contractility index) (**Fig. 7b-i**). There were no significant differences in sLVP, MAP, and heart rate between SuHx+Vehicle and SuHx+SRT2104 groups (**Fig. S8d-f**). In aggregate, these data demonstrate that SRT2104 reverses established PH and improves RV morphology and function in a severe irreversible rat model of experimental PH.

**Figure 7.**
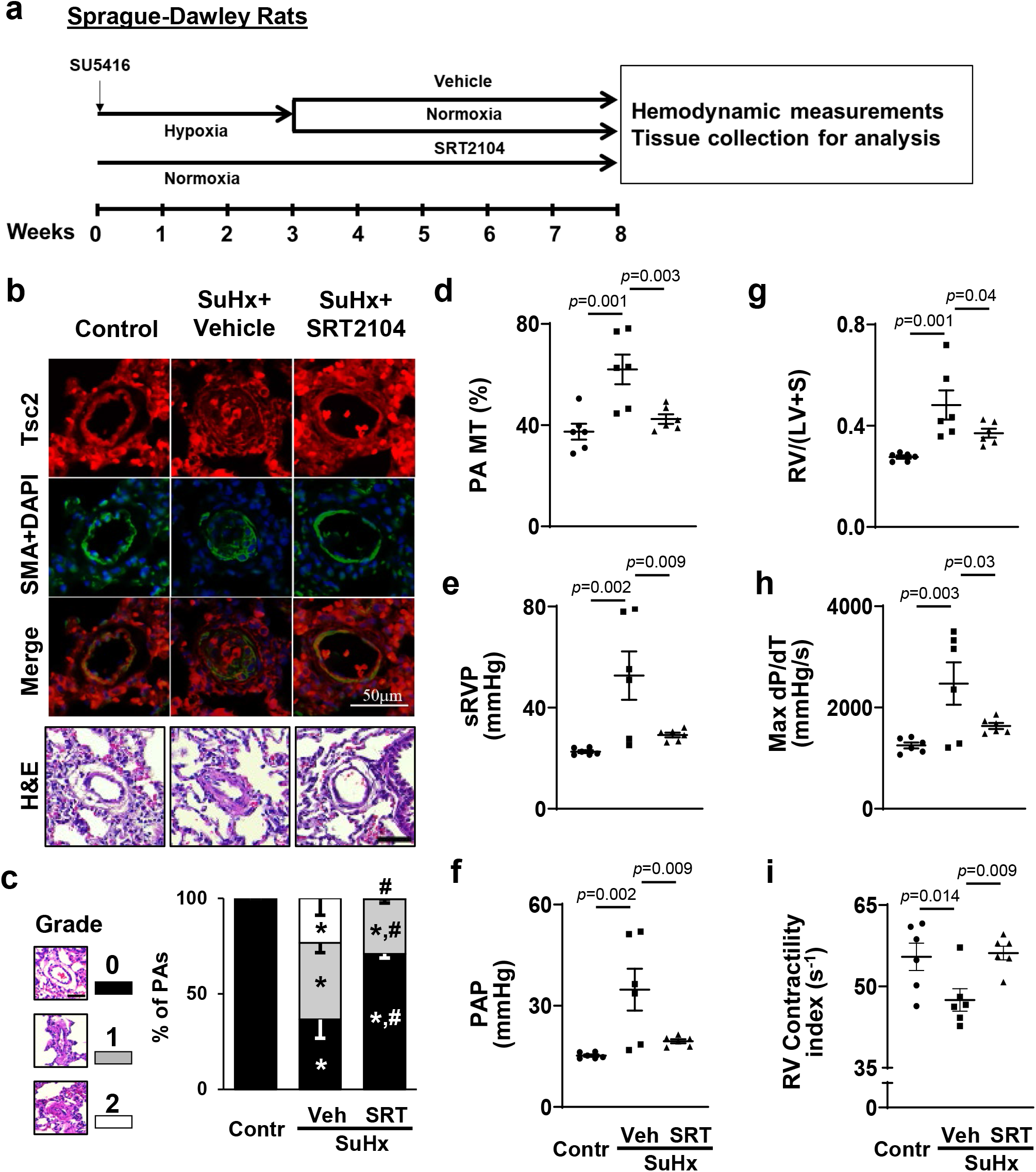
SRT2104 restores Tsc2 in small PAs, attenuates PH and reduces RV hypertrophy in rats. (see also Supplemental Fig. S8 for additional data) **a.** Schematic representation of the experiment. Male 6-8 weeks old rats received one SU5416 injection and maintained under hypoxia for three weeks. Starting week 4, rats, transferred to the normoxia conditions, were randomly assigned to two groups and then treated with SRT2104 (SRT) or vehicle (Veh) for 5 weeks, 5 days/week (days 21-56 of experiment) followed by hemodynamic and morphological analyses. Controls were same-age male rats kept under normoxia for 8 weeks. **b.** IHC to detect Tsc2 (red), SMA (green) and DAPI (blue) and H&E analyses; bar equals 50 μm, images are representative from 6 rats/group, 12 PAs/rat. **c-i**: percentage of fully (grade 2), partially (grade 1) and not occluded PAs (grade 0), bar equals 50 μm (**c**), PA MT (**d**), sRVP (**e**), PAP (**f**), Fulton index (**g**), RV contractility (Max(dP/dt)) (**h**), and RV Contractility index (**i**) measured at day 56 of experiment. Data are means±SE from 6 rats/group. \**p*<0.05 vs control, ^#^*p*<0.05 vs PH by one-way ANOVA (significance *p*<0.05).

## Discussion

The present study provides strong evidence that TSC2 deficiency in the medial layer of small PAs is an important trigger of pulmonary hypertension, and suggests that the restoration of functional TSC2 could be considered as a potentially attractive therapeutic strategy to reverse existing pulmonary vascular remodeling and PH. Our new findings are: (i) TSC2 is deficient in small remodeled PAs from subjects with PAH, two models of experimental PH, and early-passage PAVSMC derived from small PAs of PAH patients, and loss of TCS2 results in increased PAVSMC proliferation, survival, pulmonary vascular remodeling, and development of spontaneous PH; (ii) TSC2 in PAVSMC acts as a mechano-sensor and mechano-transducer and prevents up-regulation of YAP/TAZ, mTOR, and cell growth induced by pathological stiffness; (iii) TSC2 controls ECM composition, and its deficiency results in production of “diseased” ECM which up-regulates YAP/TAZ, mTOR, and induces growth of PAVSMC and PAAF; and (iv) pharmacological restoration of functional TSC2 by SRT2104 reverses molecular abnormalities, inhibits proliferation and induces apoptosis in human PAH PAVSMC, and reduces established pulmonary vascular remodeling and pulmonary hypertension *in vivo* in two models of experimental PH (**Fig. 8**).

**Figure 8.**
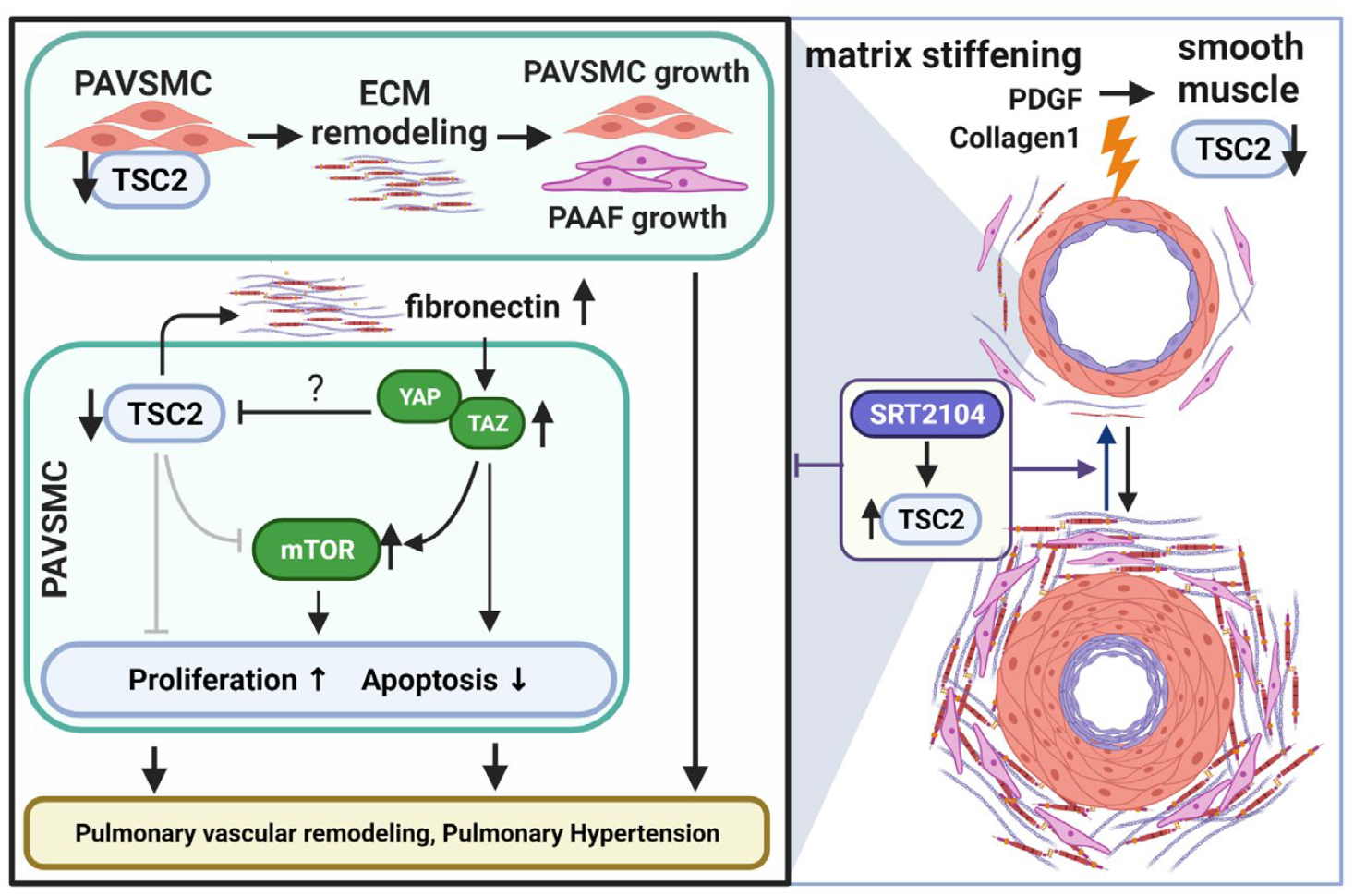
Schematic representation of the mechanism by which TSC2 loss in PAVSMC promotes pulmonary vascular remodeling and pulmonary hypertension.

Increased proliferation and resistance to apoptosis of pulmonary vascular cells in small PAs underline pulmonary vascular remodeling, a key component of PAH pathogenesis. Here, we show that TSC2, a key negative regulator of mTORC1 and cell growth (29), is deficient in proliferative PAVSMC and small remodeled PAs from human PAH lungs and in two rodent models of experimental PH. The key role of PAVSMC-specific TSC2 loss in pulmonary vascular remodeling is well supported by our *in vitro* and *in vivo* loss-of-function studies. Our data indicate that depletion of TSC2 induces unstimulated growth and proliferation of non-diseased human PAVSMC, and SMC-specific Tsc2 depletion in mice results in pulmonary vascular remodeling and early onset of spontaneous PH, indicating that even partial depletion of smooth muscle Tsc2 is sufficient to induce pulmonary vascular remodeling and pulmonary hypertension.

A switch to the hyper-proliferative, apoptosis-resistant PAVSMC phenotype could be induced by various pro-PH factors, including exposure to soluble mitogens and changes in the ECM composition and stiffness (9, 30, 31). Intriguingly, we found that TSC2 in PAVSMC is predominantly regulated by the matrix stiffness and that TSC2 deficiency, induced by pathological matrix stiffening, is required for stiffness-induced growth of PAVSMC, suggesting that TSC2 acts as a mechanosensor and a mechanotransducer. Interestingly, in addition to the pathological stiffness, TSC2 was down-regulated, although to a lesser extent, by its canonical upstream inhibitor PDGF (29), suggestive of the multi-factorial nature of TSC2 regulation. Importantly, we found that TSC2 loss was required for the accumulation of pro-proliferative/pro-survival mechanotransducers YAP/TAZ, implicated in the pathogenesis of PAH (8, 12, 32) and up-regulation of Akt-mTOR axis, indicating that TSC2 integrates mTOR and HIPPO networks, two major regulators of pulmonary vascular cell proliferation and survival in PAH (7–9, 33). Further, TSC2 is a well-described regulator of cellular metabolism, protein translation and autophagy, and loss or mutational inactivation of TSC2 in pulmonary LAM activates growth-promoting mTORC1, and up-regulates RhoA GTPase and CDK2 (34), the pro-proliferative and vasoconstrictive molecules involved in PAH pathogenesis (35, 36). Although further studies are needed, it is possible that TSC2 in PAH acts as an integrator of mechanobiological cues with PAVSMC contractility, cellular metabolism, proliferative signaling and cell cycle progression. If this is the case, the TSC2 deficiency, observed by us in “diseased” PAVSMC, could explain the complex molecular reprograming seen in PAH pulmonary vasculature and may be responsible for both, vasoconstriction and pulmonary vascular remodeling, two major pathological features of PAH.

The proposed central role of TSC2 in pulmonary vascular remodeling is further supported by our findings that TSC2 modulates ECM production by PAVSMC. We demonstrate that TSC2 deficiency in PAVSMC leads to over-production of two major ECM components, fibronectin and collagen 1 (8, 37) and formation of the “diseased” ECM with PAH-specific properties. It is also important to note that decellularized matrix, produced by control PAVSMC with shRNA-induced TSC2 depletion, was able to induce growth of not only PAVSMC, but also PA adventitial fibroblasts, demonstrating a new role of TSC2 in modulating cell-cell communications in the pulmonary vasculature. Our findings are in good agreement with evidence highlighting the importance of cell-matrix and cell-cell communications in PAH pathogenesis (37) and suggest that TSC2 could be considered as an attractive molecular target to disrupt mechanotransduction via YAP and inhibit ECM remodeling and the associated proliferation of pulmonary vascular cells in PAH.

In the TSC2-deficient perivascular epithelioid tumors, YAP accumulation is induced by mTOR via impaired autophagosomal/lysosomal degradation (21). Our findings suggest the novel mechanism of TSC2-dependent YAP/TAZ and mTOR regulation via extracellular matrix and provide the link from TSC2 via extracellular fibronectin and integrin α5β1 to the PAVSMC growth response. Further, our observations that YAP negatively regulates TSC2 in an ECM-independent way provide evidence of the existence of negative intracellular TSC2-YAP cross-talk that supports unstimulated self-sustained proliferation and apoptosis resistance of PAVSMC in PAH.

Our study identifies TSC2 as an attractive molecular target, and SIRT1 activator SRT2104 as a potential therapeutic approach to restore TSC2 and reverse pulmonary vascular remodeling and pulmonary hypertension. Reconstitution of TSC2 by mammalian vector-induced expression fully normalized PAH-specific molecular abnormalities, ECM production, reduced abnormal growth and proliferation and induced apoptosis in human PAH PAVSMC. Similar effects were observed in human PAH PAVSMC treated with SIRT1 activator SRT2104. SIRT1 plays a protective role in systemic vasculature, reducing vascular ageing and atherosclerosis (38), and its activation improves systemic arterial vascular stiffness in smokers and subjects with type 2 diabetes. Mechanistically, SIRT1 stabilizes TSC2 via lysine deacetylation (24), preventing its degradation. We show that SRT2104 has a similar ability to reverse the diseased PAH PAVSMC phenotype, reduce proliferation and induce apoptosis via restoration of TSC2. Our *in vivo* data further support the potential attractiveness of SRT2104 as a therapeutic approach to treat PH. Orally administered SRT2104 restored smooth muscle Tsc2 in small PAs in two animal models of SuHx-induced PH. Importantly, even short-time (seven days) treatment of mice with already developed PH led to reduced pulmonary vascular remodeling, sRVP, and RV hypertrophy, the central clinical features of this disease. Further, longer treatment of rats with already established PH reversed pulmonary vascular remodeling, sRVP, Max dP/dT, and RV hypertrophy, and improved the RV contractility index, making TSC2 a promising target and SRT2104 a promising drug for therapeutic intervention.

In agreement with our findings, a polyphenol resveratrol which acts, at least in part, via activating SIRT1, prevented monocrotaline-induced PH in rats (39). Resveratrol, however, is extensively metabolized in humans, resulting in low systemic bioavailability and limited potential for therapeutic intervention (40). A rapamycin-based mTORC1 inhibitor, ABI009, is now in a clinical trial for patients with severe PAH (ClinicalTrials.gov Identifier: NCT02587325) and shows promising interim results (41); however, intravenous delivery and reported side effects could make use of this drug challenging for PAH patients. SIRT1 activator SRT2104 has an improved bioavailability, demonstrates benefits in pre-clinical models of age-related disorders (42), good tolerability in humans, a favorable selectivity profile (43), and has already entered clinical trials for type 2 diabetes and psoriasis (44, 45), which makes it an attractive candidate for clinical trials for patients with PAH.

We recognize that our study has several limitations. (1) One of the limitations is a small human sample size that arises from the nature of the studied disease. PAH is rare disease, which limits the availability of human lung tissue specimens and cells of early passage for mechanistic research of this type. However, the causal role of TSC2 deficiency in mediating PAVSMC hyper-proliferation and pulmonary vascular remodeling in PAH is supported by the siRNA-mediated TSC2 depletion studies and studies of transgenic mice with SM-specific Tsc2 deficiency. (2) Not all *in vitro* experiments are designed using hydrogel matrices to model physiological or pathological stiffness. To separate stiffness-dependent TSC2 functions from its role in ECM remodeling and self-sustained proliferation and survival of PAH PAVSMC, several sets of data were collected from the cells cultured on plastic. While in agreement with our human tissue-based and hydrogels-derived data, the different substrate may affect a magnitude of observed molecular changes. (3) Because we didn’t observe any differences between males and females in the SRT2104-treated SuHx mice, only male rats were evaluated in the rat SuHx model of PH. Finally, (4) rodent models of PH, can’t completely recapitulate human PAH; the SRT2104 was given to the animals as a mono-treatment, and the combination of SRT2104 with currently used FDA-approved therapies is not evaluated. However, given our current findings in the cells from PAH subjects and transgenic mice, further testing of targeting TSC2 to reverse PAH is worthy of further investigation.

Despite the major progress in PAH treatment within the past decade, there are still no curative options to reverse advanced disease. This is largely attributable to the multifactorial nature of this disease and complex molecular and metabolic reprograming of pulmonary vascular cells supported by cell-cell and cell-matrix interactions. Our finding that TSC2 links mechanobiological cues, growth factors and ECM signals with hyper-proliferation of PAVSMC and PAAF offers new options for therapeutic intervention. Our preclinical evidence show that SRT2104, which is already in clinical trials for other diseases and has a favorable safety profile (46), shows beneficial effects in human PAH PAVSMC and two rodent models of PH, warranting further assessment in other pre-clinical models of PH and clinical trials.

## Materials and Methods

### Human Tissues and Cell Cultures

Human lung tissues from unused donor (control) and idiopathic PAH patients were provided by the University of Pittsburgh Medical Center Lung Transplant, Pulmonary Division Tissue Donation under protocols approved by the University of Pittsburgh institutional review boards. Human primary distal PAVSMC, primary pulmonary adventitial fibroblasts (PAAF) and primary pulmonary arterial endothelial cells (PAEC) were provided by the Pulmonary Hypertension Breakthrough Initiative (PHBI) or by the University of Pittsburgh Vascular Medicine Institute (VMI) Cell Processing Core. Cell isolation, characterization and maintenance were performed under the rigorous and well-established study protocols adopted by the PHBI as described in (7, 8). Briefly, pre-capillary PAs were dissected out from the left lower lobe and adherent lung parenchymal tissue using microscissors and scalpel. The arteries were minced to 1mm^2^ blocks, placed on the tissue culture plates with a small drop of LONZA culture medium (LONZA, Walkersville, MD) supplemented with SmGM-2 or FGM-2 media kit for PAVSMC and PAAF isolation, respectively (7, 8). On the following day, a full volume of respective full growth culture medium was added to the plates and left undisturbed for 3-5 days. The medium was then subsequently changed every other day until the cells reached confluence. PAVSMCs were characterized using antibodies against three smooth muscle-specific markers (smooth muscle α-actin, smooth muscle myosin heavy chain, and SM22; Cell Signaling Technology, Danvers, MA) and cell morphology (7, 8). PAAFs were characterized using antibodies against vimentin, CD90, and negative staining for SM22, vWF, and cytokeratin (Cell Signaling Technology, Danvers, MA), to exclude contamination with smooth muscle, endothelial and epithelial cells, respectively (47). Primary cells (3-8 passage) of the same passage from a minimum of three control and three PAH subjects were used for each experimental condition. Cells and tissues from de-identified human subjects were used (please see Supplemental **Table S1** for human subjects characteristics). All cells were maintained at 37 °C in a humidified incubator with 5% CO_2_. For serum-deprivation, cells were maintained for 24-48hr in basal PromoCell medium (PromoCell, Heidelberg, Germany) supplemented with 0.1% bovine serum albumin (BSA) (Thermo Fisher Scientific, Waltham, MA). Softwell hydrogel-coated plates were purchased from Matrigen (Brea, CA) and used following the manufacturer’s protocol.

**Apoptosis analysis** was performed using the *In Situ* Cell Death Detection Kit (Roche, Nutley, NJ) based on terminal deoxynucleotidyltransferase-mediated dUTP-biotin nick end labeling (TUNEL) technology according to the manufacturer’s protocol as previously described in (7, 8, 48). Nuclei were detected by 4′,6-diamidino-2-phenylindole (DAPI) (Invitrogen, Carlsbad, CA).

Images were captured by an All-in-One Fluorescence Microscope BZ-X810 (Keyence, Itasca, IL); blind semi-automatic counts were performed. A minimum of 200 cells were counted per each condition in each experiment.

**Cell proliferation** was examined using DNA synthesis analysis (bromodeoxyuridine (BrdU) incorporation assay) or Ki67 detection with specific antibody (Cell Signaling Technology, Danvers, MA) as described previously (7, 8, 48, 49). For the BrdU incorporation assay, cells, serum-deprived for 48 hours, were incubated with 10μM BrdU (Abcam, Cambridge, MA) for 18 hours, fixed with 4% paraformaldehyde in PBS (Santa Cruz Biotechnology, Dallas, TX), permeabilized by 2M HCl, and stained with anti-BrdU antibody (BD Biosciences, San Jose, CA). Staining with DAPI and anti-GFP antibody (Cell Signaling Technology, Danvers, MA) was performed to detect nuclei and GFP, respectively. For Ki67 detection, staining with anti-Ki67 antibody and DAPI was performed. Images were taken using an All-in-One Fluorescence Microscope BZ-X810 (Keyence, Itasca, IL); blinded semi-automatic analysis was performed. The cell proliferation was evaluated as the percentage of BrdU-positive or Ki67-positive cells per total number of cells. A minimum of 200 cells were counted in each experiment per each condition.

**Cell growth analysis** was performed as described previously in (7, 8, 48). Briefly, cells, treated with ATN-161 (MCE LLC, Monmouth Junction, NJ), BTT-3033 (R&D Systems, Minneapolis, MN), SRT2104 (InvivoChem, Libertyville, IL), or an appropriate diluent were trypsinized and resuspended in the same volume of basal medium supplemented with 0.1% BSA. Cell counts were performed using the Countess II FL Automated Cell Counter (Thermo Fisher Scientific, Waltham, MA).

**Immunohistochemical, Immunocytochemical and Immunoblot Analyses** were performed as described in (7, 8, 19, 50). Images were captured with an Olympus FV1000 Confocal Microscope (Olympus America Inc., Central Valley, PA) or the All-in-One Fluorescence Microscope BZ-X810 (Keyence, Itasca, IL). Immunoblot signals were captured with HyBlot CL® Autoradiography Films (Thomas Scientific, Swedesboro, NJ), and fixed and developed by Medical Film Processor (Konica Minolta, Ramsey, NJ). Immunoblots and immunostainings OD were analyzed using ImageJ (NIH, Bethesda, MD). Antibodies for TSC2 (#4308), Cleaved Caspase-3 (Asp175) (#9661), GFP (#2956), phospho-Ser235/236-S6 (#4856), S6 (#2217), YAP/TAZ (#8418), α/β-Tubulin (#2148), phospho-Ser473-Akt (#4060), Akt (#9272), and anti-rabbit IgG-HRP-linked antibody (#7074) were purchased from Cell Signaling Technology (Danvers, MA); anti-fibronectin antibody (ab2413) was purchased from Abcam (Cambridge, MA); anti-collagen1A1 antibody (AF6220) was purchased from R&D Systems (Minneapolis, MN); anti-actin, and α-smooth muscle-FITC antibody (F3777) were purchased from Sigma-Aldrich (St. Louis, MO); Alexa Fluor 488, 594 and 647 dyes and rabbit anti-sheep IgG-HRP-linked antibody (#61-8620) were purchased from Thermo Fisher Scientific (Waltham, MA).

**Transfection, Infection, and de-cellularization** were performed as described previously in (7, 8, 51). The p-EGFP plasmid (#6077-1) was purchased from Addgene (Watertown, MA); the pEGFP-TSC2 plasmid was created as previously described in (52, 53). siRNAs and shRNAs were purchased from Dharmacon (PerkinElmer, Waltham, MA) and Santa Cruz Biotechnology (Dallas, TX), respectively. siRNA transfection and shRNA infection were performed according to the manufacturers’ protocols. The Effectene transfection reagent (Qiagen, Valencia, CA) was used following the manufacturer’s protocol. For the production of decellularized matrices, pre-confluent PAVSMC were maintained on the plastic plates for six days, and then decellularization was performed.

#### Animals

All animal procedures were performed under the protocols approved by the Animal Care and Use Committees of the University of Pittsburgh and the University of California, Davis.

<u>*SM22-Cre/Tsc2^+/−^ mice*</u> were developed by crossing female Tsc2^f/f^ mice (Tsc2^tm1.1Mjg^/J, #027458) with male SM22-Cre mice (B6.Cg-Tg(Tagln-cre)1Her/J, #017491) (Jackson Laboratory, Bar Harbor, ME) carrying mouse smooth muscle protein 22-alpha (SM22) promoter inducing Cre recombinase expression selectively in the VSMC (54, 55). Nine weeks-old animals were subjected to hemodynamic and morphological analysis as described below. All mice were genotyped prior to experiments. The genotyping of the DNA extracted from ears of transgenic mice by Extract-N-Amp™ Tissue PCR Kit (Sigma St. Louis, MO) was performed following the manufacturer’s protocol. Polymerase chain reaction (PCR) samples were prepared using KAPA2G Fast PCR Kit (Sigma St. Louis, MO) following the manufacturer’s protocol. *SM22-Cre* gene was detected by Cre primers (Cre-R: 5’-GCAATTTCGGCTATACGTAACAGGG; Cre-F: 5’- GCAAGAACCTGATGGACATGTTCAG) with internal control primers (oIMR7338: 5’- CTAGGCCACAGAATTGAAAGATCT; oIMR7339: 5’-GTAGGTGGAAATTCTAGCATCC). *Tsc2* gene was detected by Tsc2 primers (Tsc2-F: 5’-ACAATGGGAGGCACATTACC; Tsc2-R: 5’-AAGCAGCAGGTCTGCAGTG). All primers were purchased from Sigma (St. Louis, MO). Controls were same-age same-sex mice of the same background (C57BL/6J, Jackson Laboratories, Bar Harbor, ME) housed in the same conditions (8, 56, 57).

<u>*In vivo experimental PH*</u> was induced by SU5416/hypoxia as described previously in (7, 8, 49). <u>*Mice:*</u> Six- to eight-week-old male and female C57BL/6J mice were exposed to hypoxia (10% O_2_) up to 35 days; subcutaneous injections of SU5416 (20 mg/kg) (Tocris, Minneapolis, MN) were performed at days 0, 7, and 14 of the experiment. At days 15-21 after PH induction, SRT2104 (InvivoChem, Libertyville, IL) suspended in the warm corn oil (ACROS Organics, Fair Lawn, NJ) (100mg/kg/day, oral gavage, 200μL/mouse) or warm corn oil alone were administrated daily. Both male and female mice were used; controls were same-age same-sex mice maintained under normoxia. <u>*Rats:*</u> Six- to eight-week-old male Sprague-Dawley rats (Charles River Laboratories, Wilmington, MA) received a single dose of SU5416 (sq 20 mg/kg) and were then maintained for three weeks under hypoxia (10% O_2_) and for five weeks under normoxia. SRT2104 (InvivoChem, Libertyville, IL) was (ACROS Organics, Fair Lawn, NJ); SRT2104 and vehicle (warm corn oil alone) were administrated starting at the beginning of the week four of the experiment for five weeks (100mg/kg/day, oral gavage, five days/week). Negative controls were normoxia-maintained age-matched animals. The volume of the vehicle was equal for the volume of SRT2104 suspension (200μL/mouse or 3mL/rat). After terminal hemodynamic analysis, animals were euthanized, and lung and heart tissues were collected for morphological analysis. Hearts were separated into right ventricle (RV) and left ventricle (LV)+Septum, and the Fulton index was calculated as RV/(LV+Septum) ratio.

<u>*Hemodynamic analysis*</u> was performed as described in (8, 49, 57, 58). Briefly, animals were anesthetized by isoflurane (Minrad Inc., Orchard Park, NY) (5% for induction, 2% during surgery, 1% while performing PV loop measurements), and *in vivo* pressure-volume PV loop measurements were performed by PV catheters (Scisense, Inc., London, ON, Canada). The catheter, attached to the data acquisition system (EMKA Instruments, Falls Church, VA) was inserted into the RV and then into the LV; data were acquired by the ADVantage PV System (Transonic Systems Inc., Ithaca, NY) and IOX2 software (EMKA Technologies Inc., Falls Church, VA). PAP was calculated as a sRVP×0.65+0.55 mmHg (49, 59). MAP was calculated as sLVP×0.65+0.55 mmHg (49, 59); RV contractility index was calculated as (Max dP/dT)/sRVP s^−1^ (49, 60).

<u>*Morphological and immunohistological analyses*</u> were performed as described previously in (7, 8, 19, 48, 49, 58, 61). Briefly, after being perfused by PBS via PA, lungs were filled with 80% Tissue Plus OCT Compound (Fisher Scientific, Hampton, NH)/20% saline solution, snap frozen in dry ice and kept at −80°C, or fixed in 4% paraformaldehyde solution in PBS and embedded in paraffin. After sectioning, lung tissue slides (5μm thickness) were stained by H&E or immunostained to detect Tsc2, smooth muscle α-actin (SMA) and nuclei (DAPI) (Invitrogen, Carlsbad, CA) as previously described in (7, 8). Images for PA MT calculation were captured from blind-selected small PAs (25-100μm outer diameter; a minimum of six animals/group, a minimum of 12 PAs/animal) of H&E- or SMA-stained rodent’s lung tissues and calculated using the VMI Calculator (61). The whole H&E-stained rat’s lung tissue sections were captured (6 rats/group, minimum of 29 PAs/rat) to count percentage of fully (grade 2), partially (grade 1), and non-occluded (grade 0) small (25-50μm outer diameter) PAs (8, 58, 61–63). Images were taken with an All-in-One Fluorescence Microscope BZ-X810.

**Statistical Analysis** was performed using STATA software (StataCorp, College Station, TX) software. Statistical comparisons between two groups were performed by the Mann-Whitney U test (64) (significance p-value less than 0.05); *in vitro* data statistical comparisons among 3 or more groups were performed by the Kruskal-Wallis test with Dunn’s Pairwise Comparison (65, 66) (significance p-value less than 0.025); *in vivo* data statistical comparisons among 3 groups were performed by the Analysis of Variance (ANOVA) test; the differences between PH or control group and other groups were analyzed by planned comparisons (67) (significance *p*-value less than 0.05).

## Non-standard Abbreviations and Acronyms

BrdU: Bromodeoxyuridine
BSA: Bovine serum albumin
Col1A1: Collagen 1A1
DAPI: 4’,6-diamidino-2-phenylindole
ECM: Extracellular matrix
ET-1: Endotelin-1
FN: Fibronectin
GFP: Green fluorescent protein
H&E: Hematoxylin and eosin
IGF-1: Insulin growth factor 1
IL-6: Interleukin-6
IPAH: Idiopathic pulmonary arterial hypertension
LAM: Lymphangioleiomyomatosis
LV: Left ventricular/ventricle
MAP: Mean arterial pressure
mTOR: Mechanistic target of rapamycin
mTORC: Mechanistic target of rapamycin complex
PA: Pulmonary artery
PA MT: Pulmonary artery medial thickness
PAAF: Pulmonary artery adventitial fibroblast
PAEC: Pulmonary artery endothelial cell
PAH: Pulmonary arterial hypertension
PAP: PA pressure
PAVSMC: Pulmonary artery vascular smooth muscle cells
PDGF-BB: Platelet-derived growth factor-BB
PH: Pulmonary hypertension
RV: Right ventricular/ventricle
SIRT1: Sirtuin 1
sLVP: Systolic left ventricular pressure
SM22: Smooth muscle protein 22-alpha
SMA: Smooth muscle a-actin
sRVP: Systolic right ventricular pressure
SuHx: SU5416/Hypoxia
TNF-α: Tumor necrosis factor-a
TSC2: Tuberous sclerosis complex 2
TUNEL: Terminal deoxynucleotidyltransferase-mediated dUTP-biotin nick end labeling
WT: Wild type
YAP: Yes-associated protein

## Supplementary Materials

Fig. S1. TSC2 protein levels are not reduced in PAAF or PAEC from PAH lungs compared to controls.

Fig. S2. Tcs2 protein reduction in SMA-positive areas in small PAs from SM22-Tsc^+/−^ mice.

Fig. S3. ECM produced by human PAH PAVSMC promotes YAP/TAZ accumulation, increases phosphorylation of Akt and S6 in control PAVSMC.

Fig. S4. YAP depletion in human PAH PAVSMC results in TSC2 accumulation.

Fig. S5. Pharmacological inhibition of mTOR has no significant effect on TSC2 and YAP/TAZ protein levels in human PAH PAVSMC.

Fig. S6. (a) TSC2 reduces Collagen 1 levels in human PAH PAVSMC. (b) BrdU incorporation in human PAH PAVSMC expressing GFP and GFP-TSC2.

Fig. S7. TSC2 protein levels in human PAH PAVSMC treated with the Akt inhibitor and AICAR.

Fig. S8. SRT2104 does not affect systemic pressure and heart rate of mice and rats with SuHx-induced PH.

Table S1. Human subjects’ characteristics.

## Acknowledgements

We thank Andrea Sebastiani and the Small Animal Hemodynamic Core at the University of Pittsburgh Heart, Lung, and Blood Vascular Medicine Institute (VMI) for acquisition and analysis of animal hemodynamic data. We thank Pulmonary Hypertension Breakthrough Initiative (PHBI) and the University of Pittsburgh VMI Cell processing Core for providing lung tissues and pulmonary vascular cells from patients with PAH and unused donor lungs. We thank John Sembrat from the University of Pittsburgh School of Medicine for his help with acquisition of human tissues.

## Funding

This work is supported by NIH/NHLBI R01HL113178 (EAG), R01HL130261 (EAG), R01HL150638 (EAG), 5P01HL103455-05 (ALM, EAG), 1 U01HL145550 (MR), 5 P50AR060780 (MR), and American Heart Association Postdoctoral Fellowship 826806 (YS). The Pulmonary Hypertension Breakthrough 13 Initiative is supported by NIH/NHLBI R24HL123767.

## Author contributions

Y.S., D.A.G., A.P., J.B., A.C.B., A.R., A.R., B.C., M.R., T.V.K., and E.A.G. acquired the data. Y.S., D.A.G., A.P., B.C., T.N.B., H.D., A.L.M., M.R., T.V.K., and E.A.G. analyzed and interpreted the data. Y.S. and E.A.G. conceived and designed research. H.D. contributed human pulmonary vascular cells. Y.S., H.D., A.M.L. and E.A.G. drafted and edited the manuscript.

## Competing interests

the authors declare no competing interests.

## Data and materials availability

The data are available from the corresponding authors on reasonable request. The materials are available via standard MTA process.

## Notes

### Competing Interest Statement

The authors have declared no competing interest.

