## Supplementary material for "TSC2-extracellular matrix crosstalk controls pulmonary vascular proliferation and pulmonary hypertension": Figures

### Supplementary Materials

**Supplemental Figures**

**Figure S1, related to Figure 1.**
**TSC2 protein levels are not reduced in PAAF or PAEC from PAH lungs compared to controls.**

**Figure S2, related to Figure 1.**
**Tcs2 protein reduction in SMA-positive areas in small PAs from SM22-Tsc$^{+/-}$ mice.**

**Figure S3, related to Figure 3.**
**ECM produced by human PAH PAVSMC promotes YAP/TAZ accumulation, increases phosphorylation of Akt and S6 in control PAVSMC.**

**Figure S4, related to Figure 3.**
**YAP depletion in human PAH PAVSMC results in TSC2 accumulation.**

**Figure S5, related to Figure 3.**
**Pharmacological inhibition of mTOR has no significant effect on TSC2 and YAP/TAZ protein levels in human PAH PAVSMC.**

**Figure S6, related to Figure 4.**
**a. TSC2 reduces Collagen 1 levels in human PAH PAVSMC.**
**b. BrdU incorporation in human PAH PAVSMC expressing GFP and GFP-TSC2.**

**Figure S7, related to Figure 5.**
**TSC2 protein levels in human PAH PAVSMC treated with the Akt inhibitor and AICAR.**

**Figure S8, related to Figures 6 and 7.**
**SRT2104 does not affect systemic pressure and heart rate of mice and rats with SuHx-induced PH.**

**Supplemental Tables**

**Table S1.**
**Human Subjects' characteristics.**

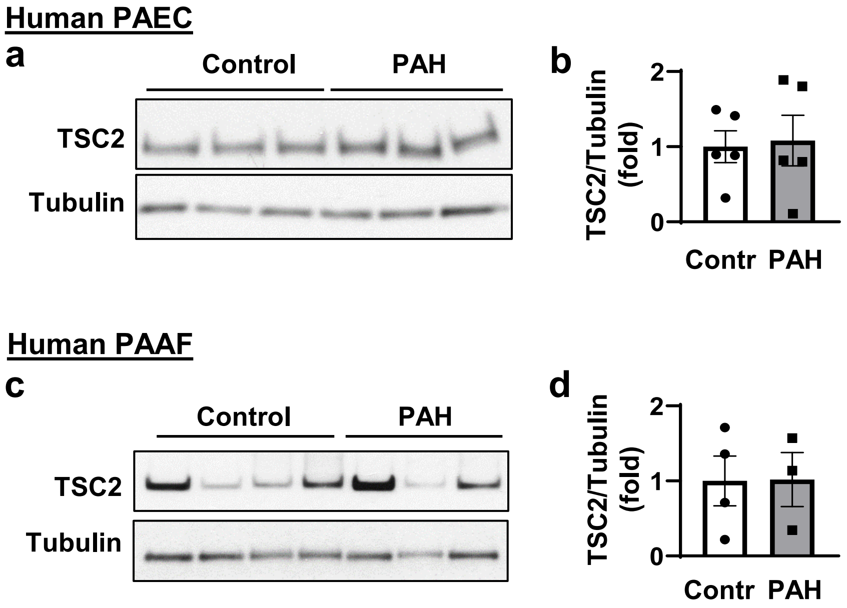


**Figure S1, related to Figure 1. TSC2 protein levels are not reduced in PAAF or PAEC from PAH lungs compared to controls**

Immunoblot analysis of PAEC (**a,b**) and PAAF (**c,d**) from non-diseased (control) and PAH subjects. Representative images (**a,c**) and analysis (**b,d**); n=5 subjects/group (PAEC), n=4 (control) and n=3 (PAH) subjects/group (PAAF). Statistical analysis by Mann Whitney U test, PAH vs. control.

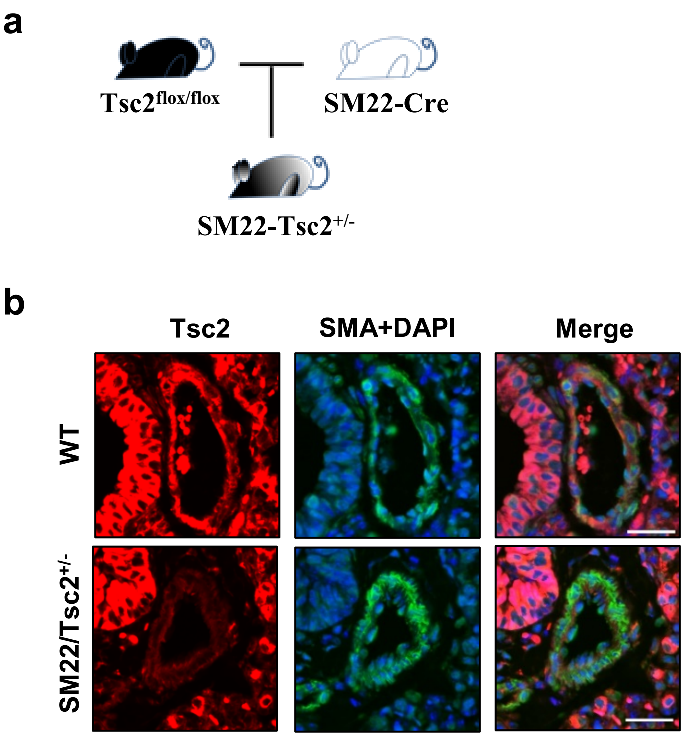


**Figure S2, related to Figure 1. Tcs2 protein reduction in SMA-positive areas in small PAs from SM22-Tsc$^{+/-}$ mice.**

**a**: Breeding scheme for developing SM22-Tsc2$^{+/-}$ mice. SM22-Cre mice were bred with Tsc2flox/flox mice to receive SM22-Tsc$^{+/-}$ mice.

**b**. Immunohistochemical analysis of lung tissue sections of SM22-Tsc2$^{+/-}$ mice to detect Tsc2 (red), smooth muscle α-actin (SMA) (green), and DAPI (blue). Bar equals 30μm; representative images from=5 (WT) and 7 (SM22/Tsc2$^{+/-}$) mice/group, respectively, 5 PAs/mouse.

#### Control PAVSMC on the matrices produced by control or PAH PAVSMC

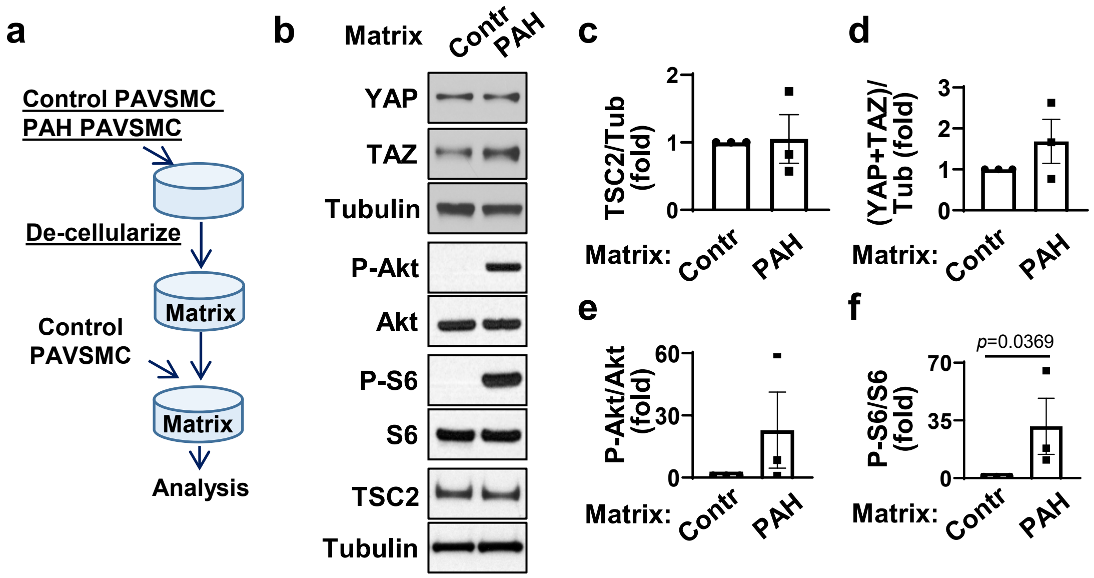


**Figure S3, related to Figure 3. ECM produced by human PAH PAVSMC promotes YAP/TAZ accumulation, increases phosphorylation of Akt and S6 in control PAVSMC**

Human PAH or control PAVSMC, reached pre-confluent condition, were grown for 7 days. Then cells were removed and equal amount of non-diseased (control) PAVSMC were plated on remaining matrices. 4 days post-plating, immunoblot analyses were performed.

**b.** Images are representative from three independent experiments, each performed on the cells from different subject.

**c-f.** data are means±SE from n=3 subjects/group, fold change to control; statistical analysis by Mann Whitney U test (significance $p<0.05$), PAH vs. control.

#### Human PAH PAVSMC

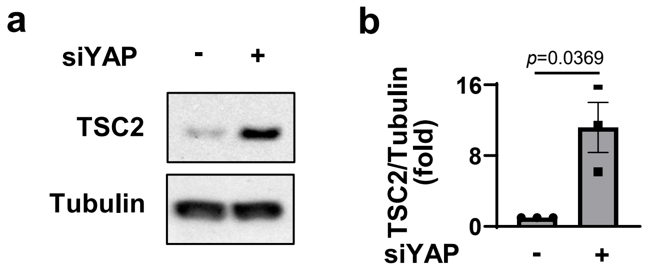


**Figure S4, related to Figure 3. YAP depletion in human PAH PAVSMC results in TSC2 accumulation.**
Immunoblot analysis of human PAH PAVSMC transfected with siRNA YAP (+) or control siRNA GLO (-) for 48hr; (a) representative images; (b) Data are means±SE, fold change to control from n=3 subjects/group. Statistical analysis by Mann-Whitney U test (significance $p<0.05$), siRNA YAP vs. siRNA GLO.

**Human PAH PAVSMC**

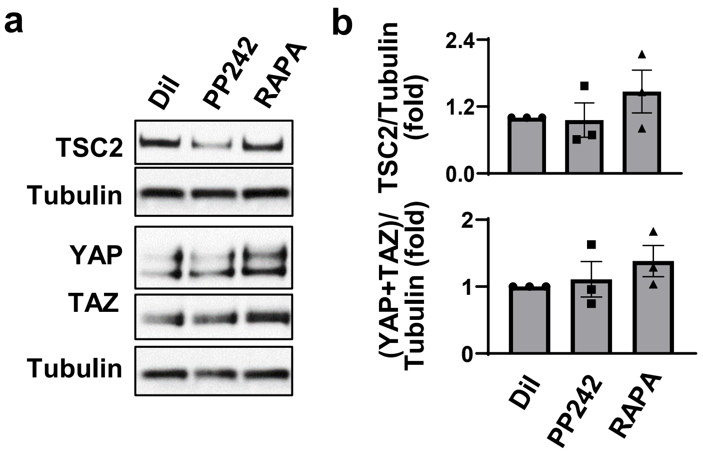


**Figure S5, related to Figure 3. Pharmacological inhibition of mTOR has no significant effect on TSC2 and YAP/TAZ protein levels in human PAH PAVSMC**

Immunoblot analysis of human PAH PAVSMC treated with 10 μM PP242, 10 nM rapamycin (RAPA), or diluent (Dil) to detect TSC2, YAP and TAZ. (a) Representative images; (b) data are means±SE, fold change to control; n=3 subjects/group. Statistical analysis by Kruskal-Wallis rank test with Dunn's Pairwise Comparison (significance $p<0.025$).

#### Human PAH PAVSMC

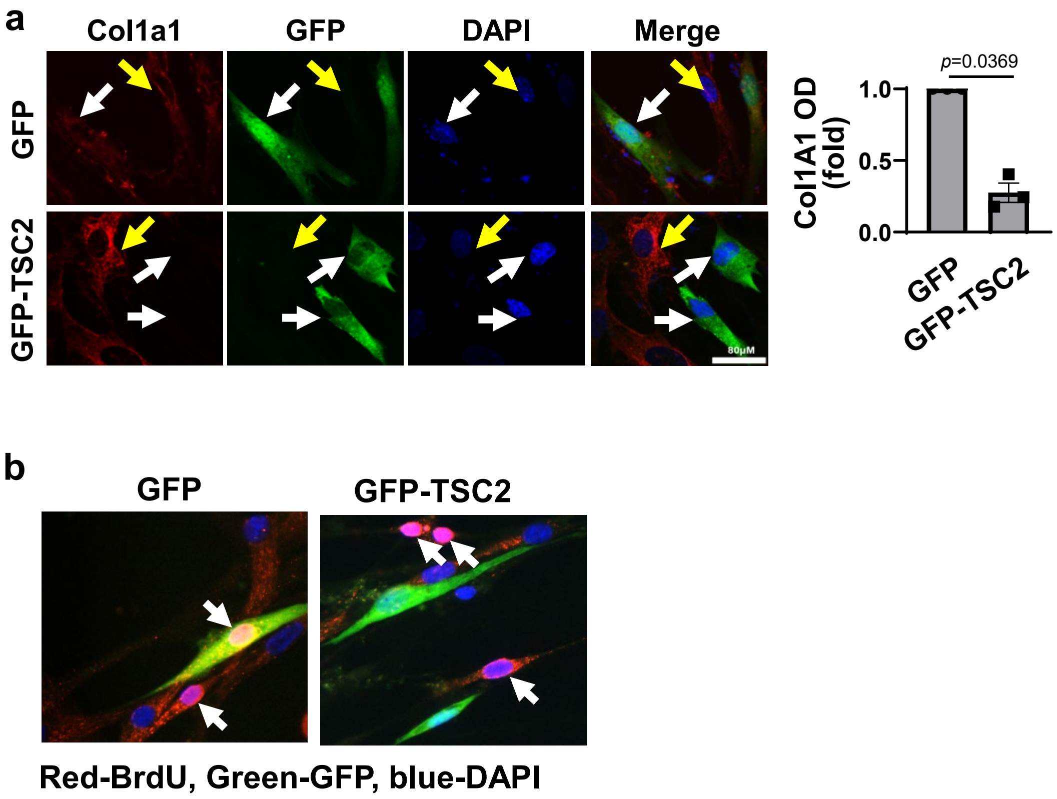


**Figure S6, related to Figure 4. TSC2 reduces Collagen 1 levels in human PAH PAVSMC**
**a.** Human PAH PAVSMC were transfected with mammalian vectors expressing GFP or GFP-TSC2. ICC analysis was performed 48hr post-transfection to detect Col1A1 (red), GFP (green) and DAPI (blue). White and yellow arrows indicate cells that were transfected or not transfected with plasmids, respectively. Bar equals to 80µm. Optical density of Col1A1 in $GFP^{+}$ cells were measured by ImageJ. n=3 subjects/group, 12 cells/subject. Data are means±SE, fold change to GFP. Statistical analysis by Mann-Whitney U test (significance $p<0.05$).
**b. BrdU incorporation in human PAH PAVSMC expressing GFP or GFP-TSC2**
Representative images of human PAH PAVSMC transfected with mammalian vectors expressing GFP (left, green) or GFP-tagged human TSC2 (right, green) and subjected to BrdU incorporation assay to detect DNA synthesis (BrdU-red, DAPI-blue). Yellow – overlap between GFP (green) and BrdU (red).

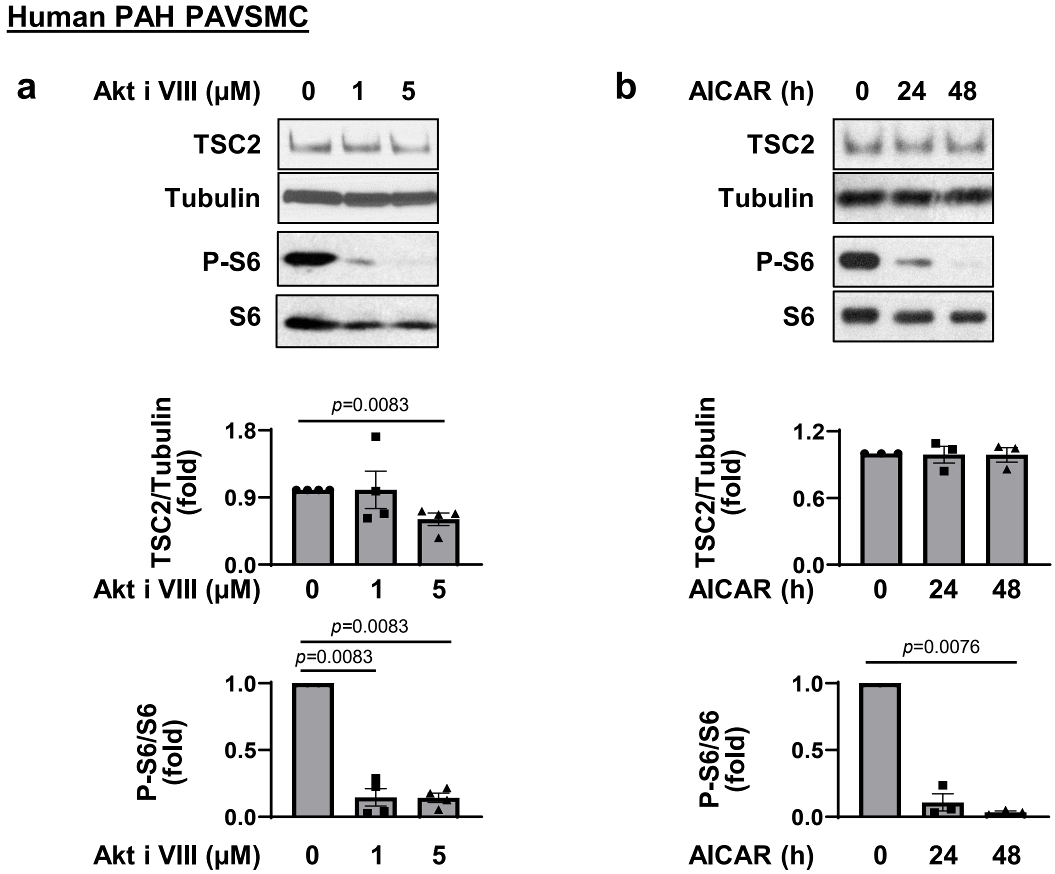


**Figure S7, related to Figure 5.**

**TSC2 protein levels in human PAH PAVSMC treated with the Akt inhibitor and AICAR**

Human PAH PAVSMC were treated with indicated concentrations of Akt inhibitor VIII for 24hr (**a**) or 100mM AMPK activator AICAR for indicated periods of time (**b**) following by immunoblot analysis to detect indicated proteins. Data are means±SE; 3 subjects/group. Significance (*p*<0.025) determined by Kruskal-Wallis test with Dunn's Pairwise Comparison vs. Diluent (0) or 0hr treatment.

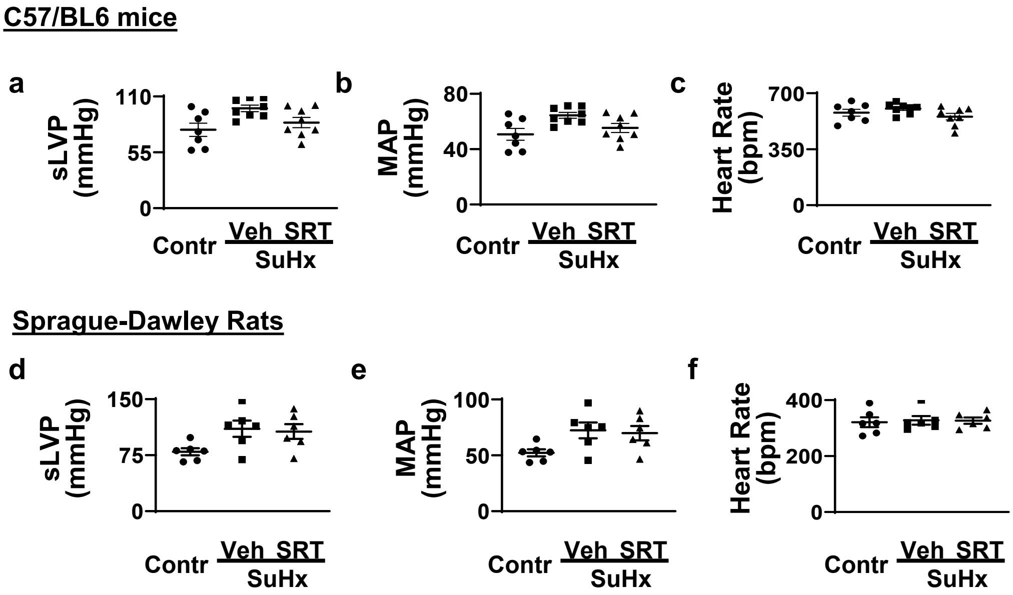


**Figure S8, related to Figures 6 and 7. SRT2104 does not affect systemic pressure and heart rate of mice and rats with SuHx-induced PH**

**a-c:** systolic left ventricular pressure (sLVP) (a), mean arterial pressure (MAP) (b), and heart rate (c) of mice without PH or with SuHx-induced PH treated with vehicle or SRT2104. Data are means±SE from n=7 mice for control (3♂ and 4♀), n=8 mice for PH+Veh (4♂ and 4♀), and n=8 for PH+SRT (4♂ and 4♀). Statistical analysis by one-way ANOVA (significance $p$<0.05).

**d-f:** sLVP (d), MAP (e), and heart rate (f) of control rats or rats with SuHx-induced PH treated with vehicle or SRT2104.. Data are means±SE from 6 rats/group. Statistical analysis by one-way ANOVA (significance $p$<0.05).

**Table 1. Human Subjects characteristics.**

| Diagnosis | Sex | Age, years |
|---|---|---|
| Non-diseased | F | 28 |
| Non-diseased | F | 29 |
| Non-diseased | F | 33 |
| Non-diseased | F | 34 |
| Non-diseased | F | 36 |
| Non-diseased | F | 37 |
| Non-diseased | F | 38 |
| Non-diseased | F | 40 |
| Non-diseased | F | 41 |
| Non-diseased | F | 43 |
| Non-diseased | F | 44 |
| Non-diseased | F | 46 |
| Non-diseased | F | 48 |
| Non-diseased | F | 50 |
| Non-diseased | F | 50 |
| Non-diseased | F | 53 |
| Non-diseased | F | 56 |
| Non-diseased | F | 57 |
| Non-diseased | F | 64 |
| Non-diseased | M | 24 |
| Non-diseased | M | 25 |
| Non-diseased | M | 35 |
| Non-diseased | M | 35 |
| Non-diseased | M | 36 |
| Non-diseased | M | 40 |
| Non-diseased | M | 41 |
| Non-diseased | M | 44 |
| Non-diseased | M | 47 |
| Non-diseased | M | 53 |
| Non-diseased | M | 70 |
| IPAH | F | 16 |
| IPAH | F | 16 |
| IPAH | F | 29 |
| IPAH | F | 32 |
| IPAH | F | 39 |
| IPAH | F | 40 |
| PAH | F | 49 |

| | | |
|---|---|---|
| IPAH | F | 50 |
| IPAH | F | 53 |
| IPAH | F | 58 |
| IPAH | F | 62 |
| IPAH | M | 21 |
| IPAH | M | 31 |
| PAH | M | 45 |
| IPAH | M | 51 |
| IPAH | M | 53 |

PAH: pulmonary arterial hypertension
IPAH: Idiopathic pulmonary arterial hypertension
